# Neural stem cells shape intracellular calcium landscapes to control cell identity and function

**DOI:** 10.64898/2026.05.20.726652

**Authors:** Bernice C. Lin, Isabella R. Maag, Hannah M. Shaw, Alessandra G. Jester, Asher B. Swan Adams, Beverly J. Piggott

## Abstract

Asymmetrically dividing neural stem cells (NSCs) provide the foundation for brain development by coupling self-renewal to the generation of diverse differentiated progeny. Yet how NSCs actively sculpt intracellular Ca²⁺ dynamics to drive developmental programs and cell behaviors across fate transitions remains poorly understood. Here we identify a role for intracellular Ca^2+^ set points in maintaining NSC identity and function in asymmetrically dividing *Drosophila* neuroblasts (NB). We find that proliferative NBs maintain low baseline cytosolic Ca^2+^, whereas differentiated progeny exhibit elevated cytosolic Ca^2+^. Experimentally increasing cytosolic Ca^2+^ slows proliferation and promotes differentiation. We further identify specific Ca^2+^ regulatory factors that are required for proliferation. Endoplasmic Reticulum (ER) luminal Ca^2+^ also differs by cell fate and depletion of ER Ca^2+^ in type II NB by loss of *SERCA* (Sarcoendoplasmic Reticulum ATPase) is sufficient to reprogram type II NB into a “type I-like” NB fate. Mechanistically, SERCA-dependent ER luminal Ca^2+^ is required for Notch receptor processing, trafficking and activation in NBs linking organellar Ca^2+^ to a core stem cell signaling pathway. Thus, NSCs and their progeny actively and distinctly shape intracellular Ca^2+^ landscapes to drive developmental programs and cell behaviors, with implications for developmental disorders and cancer.

## INTRODUCTION

Nervous systems are extraordinarily complex. Understanding how neuronal diversity arises and is precisely wired during brain formation is a fundamental question in neuroscience. Early in development, morphogen gradients pattern the neuroectoderm to establish heterogeneous populations of neural stem cells (NSCs) ^1–4^. NSCs maintain their identity while generating progressively differentiating progeny through asymmetric cell division. Asymmetrically dividing NSCs polarize cell fate determinants that unequally segregate between daughter cells during cell division so that one daughter retains self-renewal factors while the other inherits differentiation determinants^5–7^. Temporal transcription factor cascades and surrounding cellular niches further diversify neuron types according to birth order and support developmental transitions, respectively ^8–11^. As differentiation proceeds, ion channels are intrinsically upregulated to shape neural morphology, refine circuitry and ultimately establish neural excitability ^12–16^. Ion channels, exchangers and pumps are also expressed within neural precursors ^17–19^. Yet, how bioelectric signaling is tuned to support divergent cell fates and how it intersects with canonical developmental signaling pathways throughout neurogenesis remains poorly understood.

Calcium (Ca^2+^) signaling is ubiquitous and among the most extensively studied bioelectric signaling mechanisms, because it governs a wide range of physiological processes across cell types ^20–23^. Ca^2+^ acts as a second messenger whose specificity arises from tightly regulated spatial and temporal dynamics. Because of its promiscuity and potency, cytosolic Ca^2+^ concentrations are kept 10,000-fold lower than extracellular concentrations and 100 times lower than intracellular stores ^24^. Cytosolic Ca^2+^ is expelled out of the cell by proteins including Na^+^/Ca^2+^ exchangers (NCX) and PMCA (Plasma Membrane Calcium ATPase) or sequestered into the endoplasmic reticulum (ER) by SERCA (sarcoendoplasmic reticulum calcium ATPase). Increases in cytoplasmic Ca^2+^ arise from either extracellular influx through Ca^2+^-permeable channels, receptors, exchangers and transporters or by inositol 1,4,5-triphosphate (IP3) – induced Ca^2+^ release from the ER ^23^. Distinct Ca²⁺ dynamics including local transients, oscillations and global spikes are decoded by Ca²⁺-binding proteins such as calmodulin (Cam) to regulate enzyme activity, ion channel properties and transcription factor networks.

Ca^2+^ signaling has been implicated in nearly every aspect of neural development from neural induction, to proliferation, migration, differentiation and synaptogenesis ^20,25–31^. Neural cytosolic Ca^2+^ dynamics are well characterized in differentiating and mature neurons, where activity dependent Ca^2+^ influx and voltage-gated channel expression drive synaptic and circuit maturation ^22,32^. Neural progenitors comparatively, appear less active and more reliant on store operated calcium entry (SOCE), including recent work indicating that a shift away from SOCE is associated with differentiation ^33^. Far less is known about how Ca^2+^ signaling is organized in NSCs and intermediate progenitors, and whether distinct intracellular Ca^2+^ landscapes across the stem-to-progenitor continuum are functionally important for fate transitions ^32^. This gap is particularly striking given that Ca^2+^ dysregulation and mutations in Ca^2+^ regulatory proteins are implicated in numerous neurodevelopmental disorders including autism spectrum disorders ^34^, attention deficit disorder and Schizophrenia ^35–37^. Many of these research studies focus on post-mitotic neural processes such as migration and synaptogenesis ^20,31^. Disruptions to neural progenitors also cause disorders like microcephaly ^38, 39^, but the extent to which progenitor dysfunctions contribute to later onset developmental conditions remains unresolved. Ca^2+^ dynamics regulate early brain formation, including radial glial cell development in rats and humans ^18,40^, and Ca^2+^ transients have been shown to control exit from quiescence in adult NSCs in mice, with low cytosolic Ca^2+^ promoting proliferation ^41^. These findings demonstrate that like canonical developmental signaling pathways, and cell cycle regulators – intracellular Ca^2+^ dynamics play central roles in NSC regulation, but the underlying mechanisms and state-specific Ca2+ signatures are poorly defined.

In addition to cytosolic signals, the ER lumen represents a major Ca²⁺ store that is essential for protein folding, quality control and signaling ^42–45^. ER luminal Ca²⁺ levels are established by SERCA-dependent uptake and are sensed by luminal chaperones and Ca²⁺-binding proteins that regulate ER homeostasis and signaling outputs ^46–48^. Perturbations in ER Ca²⁺ can alter the unfolded protein response, modulate processing of transmembrane receptors and influence cell survival versus differentiation decisions ^49–51^, yet a causal role for ER luminal Ca²⁺ in stem cell specification *in vivo* remains largely unexplored. Whether NSCs and their progeny sculpt distinct ER Ca²⁺ landscapes during lineage progression, and whether these luminal Ca²⁺ states coordinate with canonical developmental signaling pathways to maintain cell identity, are unknown questions with broad implications for developmental disorders and disease.

Fundamental mechanisms of NSC development have been discovered using *Drosophila melanogaster* (fruit fly). Fly NSCs, known as neuroblasts (NB) divide asymmetrically to self-renew and generate more differentiated progeny, providing a powerful system in which lineage relationships, division modes and molecular identities are well defined ^6,9,52–56^. Type I NB (T1NB) reside within the brain lobes and ventral nerve cord and express the transcription factors Deadpan (Dpn) and Asense (Ase). T1NB asymmetrically divide to self-renew and generate a more differentiated Ase^+^ ganglion mother cell (GMC) that undergoes a terminal symmetric division to generate neurons or glia ^56,57^. Type II NB (T2NB) are Dpn^+^, Ase^−^. They generate an intermediate neural progenitor (INP), that once matured, expresses Ase^+^ and Dpn^+^ and divides asymmetrically to self-renew and produce GMCs^58–61 58–60^. Extensive work in *Drosophila* has established conserved cellular machinery governing asymmetric cell division and identified major regulators of identity, self-renewal and differentiation in these lineages. However, how cytosolic and ER luminal Ca^2+^ are patterned across NB, INP, and GMC states and how Ca^2+^ signaling contributes to fate transitions and lineage architecture, remain largely unexplored.

In this study, we found that cell types within *Drosophila* NB lineages establish distinct intracellular Ca^2+^ levels that are critical for maintaining cell identity and function. We discovered that NBs exhibit low cytosolic Ca^2+^ and that these Ca^2+^ levels increase in differentiated progeny. Low cytosolic Ca^2+^ supports proliferation, whereas elevated cytosolic Ca^2+^ slows proliferation and promotes differentiation. We further determined that ER luminal Ca^2+^ levels are vital for T2NB cell fate. Surprisingly, we discovered that knockdown of *SERCA*, depletes ER luminal Ca^2+^ and transforms T2NB into a “type I-like” NB identity. These transformed NBs display T1NB-specific markers and produce GMCs instead of INPs. Our data also indicates that SERCA-dependent regulation of ER luminal Ca^2+^ in the T2NB lineage is necessary for Notch signaling and maintenance of T2NB identity. Together, our findings support a model in which developing NBs intrinsically modulate intracellular Ca^2+^ dynamics to maintain cellular identity and to promote canonical developmental signaling pathways.

## RESULTS

### Ca^2+^ regulatory molecules are important for neuroblast development

To identify bioelectric regulators of neurogenesis, we used RNAi to knockdown ion channels, exchangers, pumps, and receptors in *Drosophila melanogaster* T2NB lineages (Fig. 1A). This candidate screen was performed at 96h ALH (hours after larval hatching) as it is the final stage of NB development prior to pupation, allowing us to capture defects that occur across developmental stages. A similar approach by the Bates lab was undertaken in the *Drosophila* wing disc which successfully identified numerous molecules essential for proper wing development ^62,63^. We focused on the T2NB lineages for several reasons. 1) There are only 8 T2NBs lineages per brain lobe, which made quantifying the effects of candidate knockdown on cell number more straightforward compared to the T1NB lineage which has approximately 100 NB per brain lobe. 2) T2NBs generates a transit-amplifying INP which expands the neural progeny approximately 5-fold compared to T1NB ^59,64^. And importantly, 3) T2NB lineages share patterns of division analogous to outer subventricular zone lineages in the primate cortex, providing a genetically tractable model to uncover conserved mechanisms of NSC regulation ^65–67^. We crossed *UAS* RNAi candidates or control (*luciferase RNAi*) with a T2NB driver line (*insc-GAL4; ase-GAL80*) that labeled cells within the lineage with nuclear localized GFP (GFP^NLS^). We then compared the average numbers of cells per lineage between RNAi knockdown candidates and control lineages to identify candidates that significantly increased or decreased T2NB lineage cell number. Our positive hits included four Ca^2+^ handling or signaling proteins which significantly influenced T2NB lineage cell numbers (Fig. 1B, C-F, H-I). Three of these candidates included: Na^+^/Ca^2+^ exchanger (NCX or Calx), Ca^2+^ binding protein Calmodulin (CaM), and sarcoendoplasmic Ca^2+^ ATPase (SERCA) (Fig. 1B), which all significantly reduced T2NB average cell number per lineage, suggesting a proliferation defect (Fig. 1I). Knockdown of one candidate, *Inositol 1,4,5,-trisphosphate receptor* (*Itpr*) led to a significant increase in cell number which indicated its loss enhanced proliferation (Fig. 1H-I).

**Figure 1.**
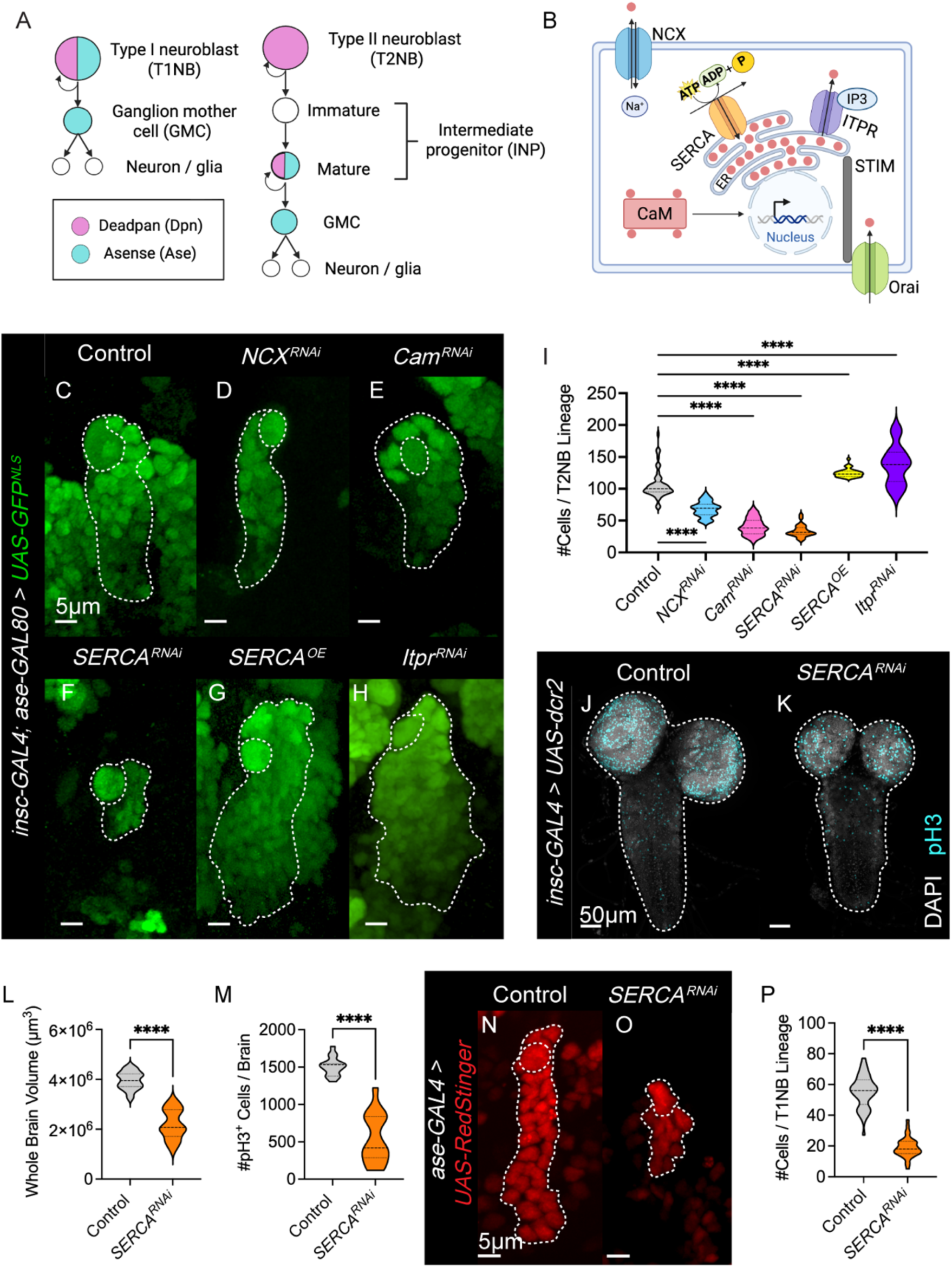
Ca^2+^ regulatory molecules are required for type II NB development. (A) Illustrative model of T1 and T2 NB lineages (B) Illustrative model of NB Ca^2+^ regulatory molecules identified from screen (C-H). Red circle = Ca^2+^, NCX = Na^+^/Ca^2+^ exchanger, CaM = Calmodulin, SERCA = Sarcoendoplasmic reticulum Ca^2+^-ATPase, ITPR = Inositol 1,4,5-triphosphate receptor, STIM = Stromal interaction molecule. (C-H) Representative images of RNAi knockdown of Ca^2+^ regulatory molecules in T2NB lineages or (G) overexpression (OE) of SERCA, labeled with nuclear localized GFP (GFP^NLS^) at 96h ALH. Control is *luciferase^RNAi^.* Dashed white lines outline NB (circle) and lineage. (I) Quantification of the number of GFP cells per T2NB lineage compared to control. One-way ANOVA with applied post hoc Dunnett’s multiple comparisons test. N ≥ 25. (J-K) Representative images of 96h ALH larval brains with *luciferase* (J) and *SERCA* (K). Dashed white lines outline larval brain. (L) Quantification of whole brain volume utilizing DAPI. Unpaired t-test. N ≥ 35. (M) Quantification of number of pH3^+^ cells in the whole brain. Unpaired t-test. N ≥ 35. (N-O) Representative images of *luciferase* (N) and SERCA (O) T1NB RNAi-knockdown in T1NB lineages at 96h ALH, labeled with RedStinger. Dashed white lines outline NB (circle) and lineage. (P) Quantification of the number of RedStinger^+^ cells within T1NB lineages. Unpaired t-test. N = 39.

As *SERCA* RNAi knockdown exhibited the strongest effect on T2NB numbers, we focused our attention on characterizing its role within NB lineages. We first verified the efficacy of the *SERCA RNAi* knockdown in NB lineages driven by *insc-GAL4* compared to a control RNAi using qPCR. We found that *SERCA* expression levels were significantly reduced upon *SERCA* knockdown (Supp. Fig. 1A). Depleting *SERCA* reduces T2NB cell numbers which suggests that the relative abundance of SERCA might influence proliferation. We next overexpressed (OE) *SERCA* in T2NB lineages, which increased cell numbers in T2NB compared to control brains (Fig. 1G, I). This indicates that SERCA abundance (and potentially resulting intracellular Ca^2+^ level changes) cell autonomously regulate proliferation. To determine if SERCA broadly influences NB populations (T1, T2, mushroom body and optic lobe NBs), we used a pan-NB driver (*insc-GAL4*) to knockdown *SERCA* and found that this significantly reduced larval brain size and the number of cells labeled with mitotic indicator, phosphohistone 3 (pH3) (Fig. 1J-M, Supp. Fig. 1B-G). To more precisely examine the effects of SERCA on T1NB lineages, we knocked down *SERCA* using a T1NB specific driver (*ase-GAL4*) and found that compared to knockdown controls, it reduced brain size (Supp. Fig. 1H”, I”, J), numbers of pH3^+^ mitotic cells (Supp. Fig.1 H-I’, K), the total number of T1NB per brain lobe (Supp. Fig1L-N), and cell numbers per individual T1NB lineage (Fig. 1N-P). These results indicate that SERCA is required for proliferation across NB lineages, demonstrating its broad necessity in brain development.

### SERCA maintains type II NB proliferation

To better understand how SERCA regulates NB lineage development we investigated the underlying basis for cell number reduction in T2NB lineages upon *SERCA* knockdown. Fewer cells within NB lineages could result from increased cell death, cell cycle defects, delayed exit from quiescence or from alterations in cellular identity. To assess if *SERCA* knockdown induced apoptosis in the T2NB lineage, we expressed baculovirus caspase inhibitor, P35, which blocks apoptotic cell death ^68,69^. P35 expression in control T2NB lineages produced a small yet significant increase in cells per lineage compared to P35-negative controls, consistent with T2NB lineages undergoing apoptosis during normal development as we and others have reported ^69,70^ (Fig. 2A-B, E). Blocking apoptosis along with *SERCA* knockdown in T2NB lineages resulted in a modest but significant increase in cell number; however, this was not sufficient to restore cell numbers to control levels (Fig. 2C-D, E). These results indicate that while *SERCA* knockdown induces apoptosis in T2NB lineages, apoptosis alone does not fully account for the observed reduction in cell number.

**Figure 2.**
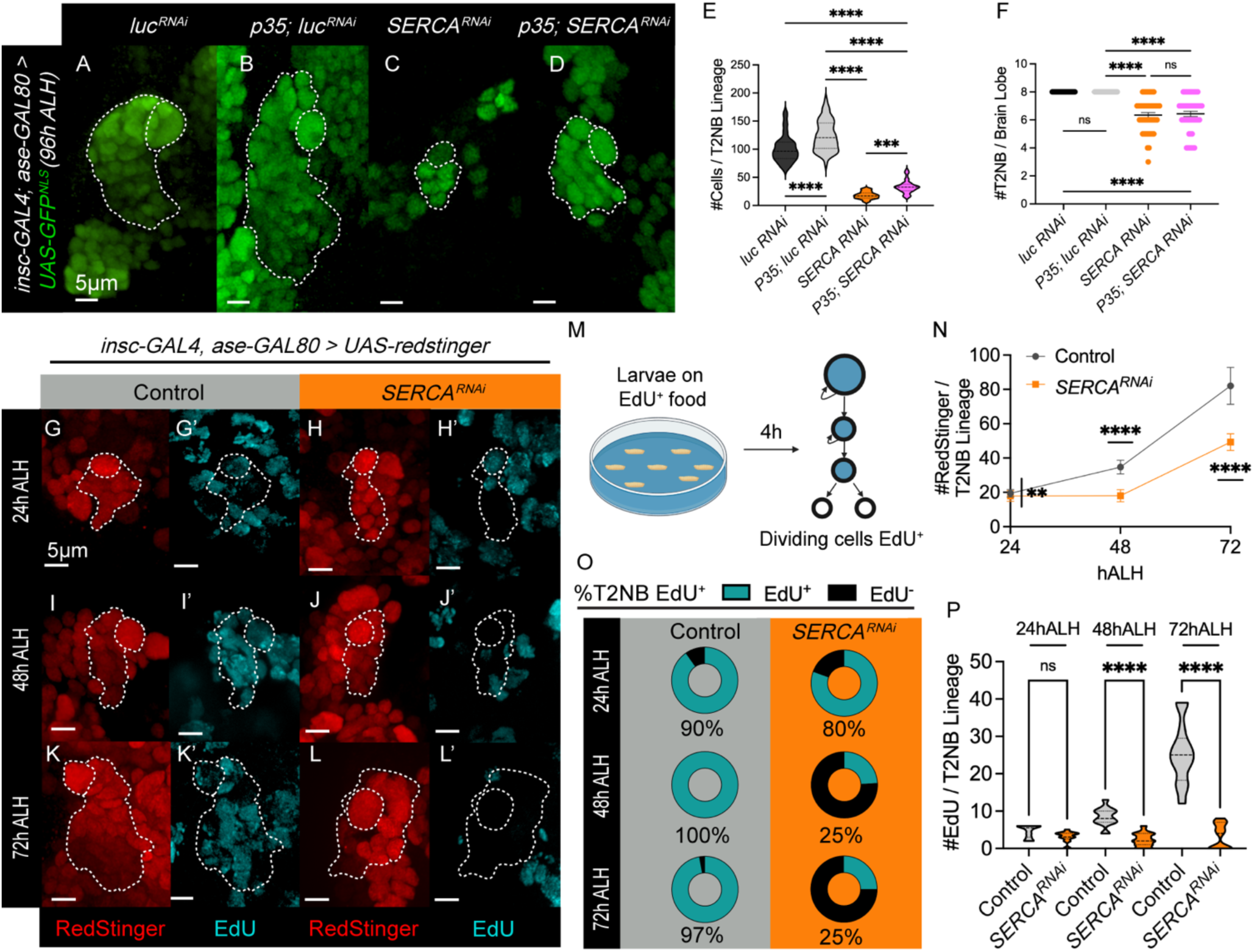
SERCA regulates type II NB proliferation and has modest effects on cell death. (A-D) Representative images of T2NB (labeled by GFP^NLS^) with and without P35 expression at 96h ALH. Control is is *luciferase^RNAi^.* Dashed white lines outline NB (circle) and lineage. (E) Quantification of (A-D) the number of GFP cells per T2NB lineage. One-way ANOVA with applied post hoc Tukey’s multiple comparisons test. N ≥ 32. (F) Quantification of number of missing T2NB per brain lobe. 96h ALH. One-way ANOVA with applied post hoc Tukey’s multiple comparisons test. N ≥ 26. Mean with SEM indicated by error bars. (G-L) Representative images of EdU staining in T2NB from 24-72hALH. (M) Illustrative model of EdU feeding experiment. All larvae placed on EdU food for 4h. (N) Quantification of the number of cells per T2NB lineage from 24-72h ALH. Nonlinear regression curve fit. N ≥ 110. (O) Quantification of the percent of T2NB that was EdU^+^. N ≥ 27. (P) Number of EdU^+^ cells per T2NB lineage. One-way ANOVA with applied post hoc Sidak’s multiple comparisons test. N ≥ 25. Dashed white lines on representative images outlines lineages. Circle white dashed line outlines NBs.

To determine if *SERCA* knockdown influences cell cycle speed, we performed an EdU (5-Ethynyl-2-deoxyuridine) incorporation assay. EdU is a thymidine analog fed to larvae during a 4-hour period that is incorporated into DNA during S-phase of mitotically active cells (Fig. 2M). There are two neurogenic periods in flies, embryonic and larval. In between these neurogenic periods, most NBs undergo quiescence—reducing their size and pausing cell division—until later reactivation in early larval stages. Once reactivated, larval NBs typically divide every 1.5h, whereas INPs and GMCs divide every 4-6h ^71^. Thus, this assay measures both how fast cells are dividing and how many NBs are still in quiescence at early larval stages. By 24h ALH most T2NBs should have exited quiescence ^72^. Quiescent NBs do not progress through the cell cycle; therefore, they do not incorporate EdU. At 24h, the majority of T2NBs from both control and *SERCA* knockdown conditions were EdU^+^ (90% EdU^+^ in control and 80% EdU^+^ *SERCA* knockdown) (Fig. 2G-H’, O). While this data indicates that *SERCA* knockdown does not universally delay T2NB exit from quiescence at early stages, a higher proportion of SERCA knockdown T2NBs failed to incorporate EdU compared to controls (20% versus 10%, respectively; Fig. 2G’-H’, O). In addition, at 24h ALH there was a slight reduction in *SERCA* knockdown NB diamaters compared to control (Supp. Fig. 2A-B, E). The number of total cells within individual T2NB lineages at 24h ALH displayed a small but significant reduction upon *SERCA* knockdown compared to control (Fig. 2N). Collectively, these results indicate that although most 24h *SERCA* knockdown T2NBs were able to progress through the cell cycle during the 4h EdU-labeling window, progenitor proliferation was modestly reduced and quiescence was partially delayed. This mild effect is unable to account for the pronounced proliferation defects observed in SERCA-deficient T2NB lineages.

At 48h ALH and 72h ALH, nearly all control T2NB lineages displayed EdU labeling (Fig. 2I-I’, K-K’, O). In contrast only about 25% of *SERCA* knockdown T2NB were EdU positive (Fig. 2I-L’, O-P). The reduction in T2NB EdU incorporation within *SERCA* knockdown larvae indicates that SERCA is required for normal cell cycle progression. Consistent with a reduction in proliferation speed, *SERCA* knockdown T2NB lineages displayed fewer cells, and fewer EdU labeled cells per lineage at nearly every stage examined (Fig. 2N, P). By 96h ALH SERCA knockdown resulted in an average loss of two of the eight T2NB per brain lobe (typically the dorsal lateral T2NB, DL1-2) (Fig. 2F). Control brains continued to maintain all eight T2NB at 96h ALH (Fig. 2F). Loss of T2NBs could reflect cell death or premature terminal differentiation. However, blocking cell death with P35 did not rescue T2NB loss in *SERCA* knockdown lineages, which indicates that the missing T2NB did not undergo apoptosis, but instead prematurely terminally differentiated (Fig. 2F). Consistent with premature cell cycle exit, 61% of *SERCA* knockdown T2NB displayed ectopic nuclear Prospero staining at 96h ALH (Supp. Fig. 2G-J). Prospero promotes neural differentiation and is normally absent from T2NB ^73^. This data aligns with previous findings that ectopic Prospero expression in T2NB leads to their premature terminal differentiation ^73^. As NBs reach the end of their proliferative period near pupation (after ∼ 96h ALH) they reduce their size and enter a phase of proliferative differentiation prior to terminal differentiation ^74^. At 72h ALH *SERCA* knockdown T2NB were significantly smaller in size than controls, consistent with their premature entry into the proliferative differentiation phase (Supp. Fig. 2C-D, F).

To investigate whether SERCA loss impacts cell cycle dynamics, we examined expression of the genetically encoded cell cycle indicator, FUCCI (Fluorescent Ubiquitination-based Cell Cycle Indicator) ^75^. Fly-FUCCI is composed of fluorescently-tagged cyclin B and E2F1 degrons whose relative expression provides a readout for cell cycle state ^75^. At 96h ALH, SERCA knockdown T2NBs shifted to G1/S phases at the expense of G2/M compared to controls (Supp. Fig. 2K-N). This indicates that a reduction in SERCA expression disrupts cell cycle progression. Despite these defects, T2NB *SERCA* RNAi lineage numbers still increased between 24h ALH to 72h ALH (Fig. 2N), demonstrating that these NB lineages retained some proliferative capacity. Together these results indicate that SERCA is required for NB proliferation and maintenance.

### Distinct intracellular Ca^2+^ dynamics promote proliferation and drive differentiation

The influence of Ca^2+^ regulatory proteins on T2NB development suggests that Ca^2+^ dynamics are important for proliferation and neural differentiation. We therefore investigated if cytosolic Ca^2+^ levels differ between the NB and their differentiated progeny within the T2NB lineage. Cell identity and differentiation state can be inferred from their morphology and position; NBs are substantially larger (∼8-10µm) than their differentiated progeny (<5µm)^71,74^. As NB divide, newly born daughter cells remain adjacent to the NB, while older more differentiated cells become progressively displaced to positions further from the NB. Thus, the further a cell is from the NB, the more differentiated it tends to be (Fig. 3H). To resolve cytosolic Ca^2+^ levels, we expressed the genetically encoded Ca^2+^ sensor, GCaMP6F, in T2NB lineages. GCaMP consists of GFP fused to a portion of CaM. It becomes brighter with increased cytosolic Ca^2+^ concentrations. We found that T2NB exhibited markedly lower GCaMP fluorescence intensity than its more differentiated progeny (Fig. 3A, C). Notably, intermediate positioned cells near the NB displayed the highest GCaMP fluorescence intensity (Fig. 3A, C). Cells furthest from the NB, likely to be neurons, showed relatively lower GCaMP intensity (Fig. 3A, C) but occasionally exhibited high Ca^2+^ transients that likely reflected neuronal activity (data not shown). Together this data indicates that there are baseline differences of cytosolic Ca^2+^ that correspond to the distinct cell fates within NB lineages (Fig. 3H).

**Figure 3.**
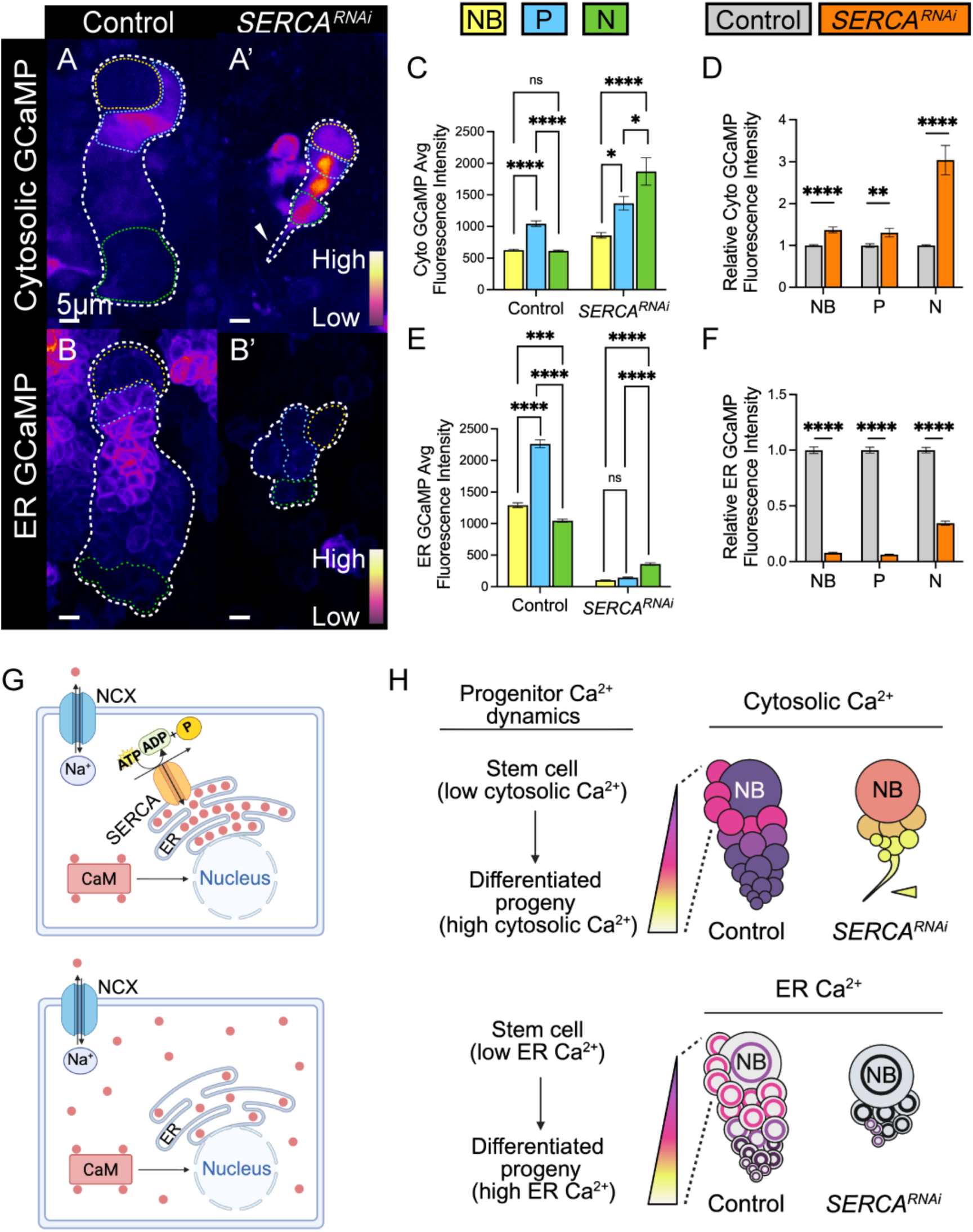
Neuroblasts exhibit distinct calcium dynamics from differentiated progeny. (A-B) Representative images of *UAS-GCaMP6f* (cytosolic, A) or *UAS-GCaMP210* (ER, B) driven in T2NB lineages with either *luc^RNAi^* (control) or *SERCA^RNAi^.* Brighter fluorescence represent higher cytosolic Ca^2+^. Arrow points to axon projection that can be visualized with *SERCA^RNAi^.* 72h ALH. Yellow dashed outline = NB. Blue dashed outline = progenitors (P). Green dashed outline = neurons (N). (C) Quantification of average cytoplasmic GCaMP fluorescence intensity of NB, P, and N in control and *SERCA^RNAi^* T2NB lineages (*insc-GAL4, ase-GAL80*) at 72hALH. One-way ANOVA with applied post hoc Tukey’s multiple comparisons test. N ≥ 53. (D) Quantification of relative cytoplasmic GCaMP fluorescence intensity between control and *SERCA^RNAi^*T2NB cell types. Unpaired t-tests. N ≥ 53. (E) Quantification of average ER GCaMP fluorescence intensity of NB, P, and N in control and *SERCA^RNAi^* T2NB lineages at 72hALH. One-way ANOVA with applied post hoc Tukey’s multiple comparisons test. N ≥ 85. (F) Quantification of relative ER GCaMP fluorescence intensity between control and *SERCA^RNAi^* T2NB cell types. Unpaired t-tests. N ≥ 85. (G) Illustrative model of intracellular changes in cytoplasmic and ER Ca^2+^ in control and *SERCA^RNAi^* T2NB lineages. Red circles = Ca^2+^. (H) Illustrative model of changes in cytosolic and ER Ca^2+^ in control and *SERCA^RNAi^*T2NB lineages. Yellow arrow points to axon projection that can be visualized with *SERCA^RNAi^*.

Because SERCA expression exerted strong effects on cell number, we investigated whether knockdown of *SERCA* influenced cytosolic Ca^2+^ levels. SERCA is localized to the ER where it helps maintain low cytosolic Ca^2+^ levels and high ER Ca^2+^ levels by using ATP hydrolysis to pump Ca^2+^ from the cytoplasm into the ER (Fig. 3G). Based on SERCA’s subcellular localization and function, reducing *SERCA* expression would be expected to elevate cytoplasmic Ca^2+^ levels. Indeed, knocking down *SERCA* in T2NB lineages at 72h ALH caused a robust increase in GCaMP fluorescence intensity across T2NB lineages, indicating an elevation of cytosolic Ca^2+^ levels (Fig. 3A-A’, C-D). These findings indicate that SERCA is required for maintaining cytosolic Ca^2+^ across cell types.

We next investigated whether elevated cytosolic Ca^2+^ was sufficient to affect proliferation independently of SERCA. To do this, we acutely elevated extracellular Ca^2+^ within *ex-vivo* brain cultures, which we reasoned would create a driving force to increase cytosolic Ca^2+^. Larval brains expressing GCaMP6F in T2NB were dissected at 96h ALH and incubated in fortified Schneiders medium (see methods) containing 5mM Ca^2+^ (control) or in Schneiders medium with 30mM (high Ca^2+^) prior to imaging (Supp. Fig. 3A-B). High extracellular Ca^2+^ significantly increased GCaMP6F fluorescence in both T2NBs and progeny compared to controls (Supp. Fig. 3C-D), indicating that elevated extracellular Ca^2+^ was sufficient to raise cytosolic Ca^2+^ levels.

Next, we tested whether elevated cytosolic Ca^2+^ affected T2NB lineage proliferation rate. Larval brains dissected at 72h ALH were either immediately fixed for lineage number analysis or incubated for 24h in control Ca^2+^ conditions or high Ca^2+^ levels (30mM) prior to fixation. Following this incubation, we fixed and stained the explant larval brains for lineage analysis. High Ca^2+^ incubation significantly reduced cell numbers per lineage (Supp. Fig. 3E-H), indicating that elevated cytosolic Ca^2+^ slows lineage expansion. These results suggest that T2NBs maintain low cytosolic Ca^2+^ levels to support robust proliferation.

In keratinocyte (skin) and osteocyte (bone) cell culture, protocols increasing extracellular Ca^2+^ levels is a well-established strategy to induce differentiation ^76–82^. Because our data indicates T2NBs display low cytosolic Ca^2+^ while differentiated progeny exhibit higher cytosolic Ca^2+^, we hypothesized that distinct cytosolic Ca^2+^ setpoints help maintain cellular identity. If this is correct, experimentally elevating cytosolic Ca^2+^ should promote differentiation. To test this idea, we examined Prospero expression in explanted brains exposed to control or high Ca^2+^ bath solutions.

High extracellular Ca^2+^ increased the proportion of Prospero^+^ cells per lineage (Supp. Fig. 3E’-G’, I), indicating that elevated Ca^2+^ promotes differentiation within T2NB lineages. Together, these results suggest a model in which NB lineages intrinsically regulate cytosolic Ca^2+^ levels to maintain cell identity: NBs keep cytosolic Ca^2+^ low to sustain proliferation, while progeny elevate cytosolic Ca^2+^ to promote differentiation (Fig. 3H). The daughter cells adjacent to the NB displayed the highest Ca^2+^ levels within the T2NB lineage (Fig. 3H), highlighting calcium’s role in the T2NB-to-daughter transition.

To determine whether ER Ca^2+^ varies with cell identity, we expressed ER-localized GCaMP (GCaMP^210^) in T2NB lineages. Like cytosolic Ca^2+^ levels, our studies revealed that ER luminal Ca^2+^ levels varied with cell identity (Fig. 3B, E). Notably, ER luminal Ca^2+^ levels were highest in cells surrounding the T2NB which are most likely to be INPs or GMCs (Fig. 3B, E, H). Reducing *SERCA* expression levels dramatically depleted ER luminal Ca^2+^ throughout the T2NB lineage (Fig. 3B-B’, E-F). The widespread reduction in ER GCaMP fluorescence intensity indicates that SERCA is the primary regulator of ER luminal Ca^2+^ levels in T2NB lineages (Fig. 3G-H). SERCA pumps Ca^2+^ from the cytosol into the ER, so one might expect that regions with high SERCA expression would exhibit low cytosolic Ca^2+^ and high ER luminal Ca^2+^. However, our data does not follow this simple prediction. In general, T2NB displayed both low cytosolic and luminal Ca^2+^ levels, whereas neural progenitors, likely INPs, exhibited higher luminal and cytosolic Ca^2+^. These differences indicate that although SERCA contributes to ER Ca^2+^ loading, additional Ca^2+^ channels, pumps and Ca^2+^-binding/buffering proteins substantially shape the free Ca^2+^ levels that we resolve with GCaMP imaging.

### Knockdown of *SERCA* transforms T2NB identity into a ‘type I like’ NB cellular fate

We found that SERCA expression levels majorly impacted the number of cells within T2NB lineages, in part, due to its influence on cell cycle progression. However, cellular identity also influences cell-cycle rate, so we next assessed which cell types were affected by reduced expression of Ca^2+^ regulatory molecules. To test this, we first investigated the effect of complete loss of *SERCA* on T2NB lineage identity using a null allele. *SERCA^S^*^5^ (*Gln108*) mutations cause early protein truncation ^45^, creating a null *SERCA* allele that is homozygous lethal but heterozygous viable. We used MARCM (Mosaic Analysis with a Repressible Cell Marker^83^) to generate homozygous *SERCA^S^*^5^ mutant clones (labeled with mCD8-GFP) in an otherwise heterozygous and viable animal^84^. Our MARCM approach utilized a one-hour heat shock which resulted in random labeling of NB lineages. In control animals, MARCM clones consistently labeled 1-2 of the 8 T2NB lineages in the third instar wandering larval stage (Fig. 4A-B’’’). T2NB are normally distinguished from T1NB by the presence of a large Dpn^+^Ase^-^ NB and the presence of small Dpn^+^ INPs that are unique to T2NB lineages. Surprisingly, we never found any MARCM-labeled T2NB lineages in *SERCA^S^*^5^ MARCM clones (Fig. 4C’’’, E). It’s possible that MARCM wasn’t labeling any T2NB lineages, however using Dpn staining, we could only resolve about 5-6 of the typical 8 T2NB lineages per lobe. Notably, the T2NBs that normally occupy the dorsal medial brain region were missing, and in their place, we found T1NB labeled homozygous *SERCA^S5^* MARCM clones. This may indicate that *SERCA^S5^* null T2NB clones were transformed into a T1NB identity or that the *SERCA^S5^* mutation is more severe than RNAi, resulting in T2NB death. To resolve this question, we returned to *SERCA* RNAi knockdown, where we could directly deplete *SERCA* in all T2NB lineages and examine the impacts on cellular identity.

**Figure 4.**
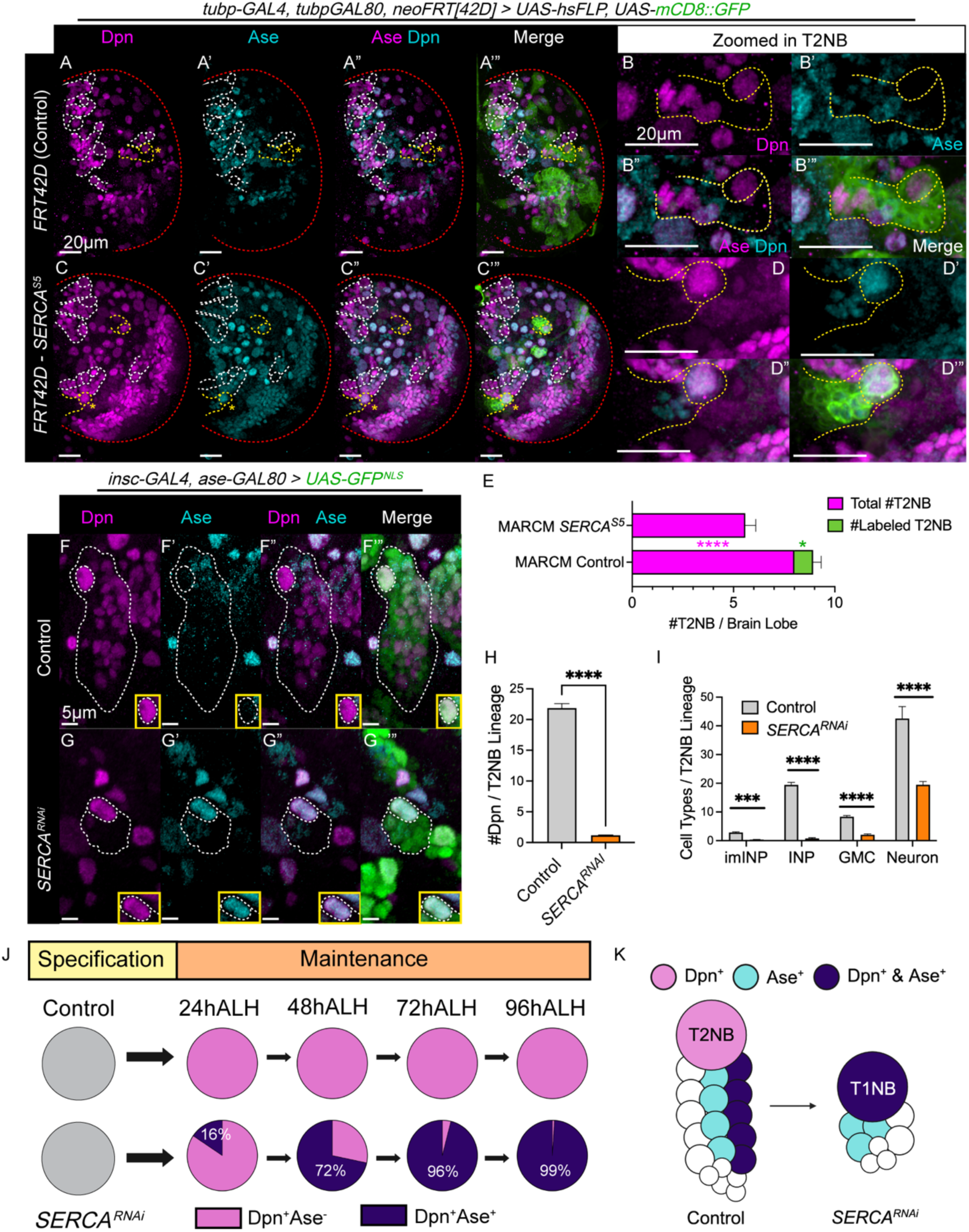
Loss of SERCA transforms type II NB into a type I-like NB identity. (A,C) Representative images of MARCM brain lobes at 96hALH with control (*FRT42D*, A) and *FRT42D-SERCA^S5^*, C). Red dashed line outlines brain lobe. White dashed lines outline unlabeled heterozygous T2NB and lineages. Yellow dashed lines outline MARCM-induced homozygous T2NB lineages labeled with mCD8::GFP. Yellow star represents the zoomed in labeled T2NB lineage in (B) and (D). (B,D) Representative images of yellow star (A) and (C) MARCM-induced homozygous T2NB labeled with mCD8::GFP. Dashed yellow lines outline NB and lineage. (E) Quantification of the total number of T2NB and MARCM-labeled T2NB per brain lobe. Unpaired t-test, N ≥ 12. (F-G) Representative images of T2NB lineages with control (*luc^RNAi^*, C), and SERCA^RNAi^ (D). Yellow inset box highlights NB. White dashed line outlines lineage and NB. 96hALH. (H) Quantification of number of Dpn^+^ cells per T2NB lineage. Independent t-test. N ≥ 42. (I) Quantification of the number of cell types per control or *SERCA^RNAi^* T2NB lineage. INP = intermediate progenitor. ImINP = immature INP. GMC = ganglion mother cell. Independent t-tests. N ≥ 23. (J) Quantification of the percentage of Dpn^+^Ase^-^ or Dpn^+^Ase^+^ NB in control and *SERCA^RNAi^* lineages from 24hALH to 96hALH. N ≥ 15. (K) Illustrative model of cell types and T2NB identity change with *SERCA^RNAi^*.

Using immunohistochemistry, we stained for Dpn and Ase which together are sufficient to identify all cell types within the T2NB lineage (Fig. 4F-F”, I) ^69^. *SERCA* knockdown significantly reduced all lineage cell types (Fig. 4I). As expected, control T2NB were Dpn^+^Ase^-^ (Fig. 4F’-F’’’); however, RNAi knockdown of *SERCA* strikingly induced ectopic expression of Ase in T2NBs (Fig. 4G’) and caused a near-complete loss of INPs (small Dpn^+^ cells) in most T2NB lineages (Fig. 4G-I). Thus, SERCA-knockdown transformed T2NBs into a ‘T1-like NB’ identity based on both marker expression and lineage composition (Fig. 4K). This data is consistent with our MARCM-labeled *SERCA^S5^* mutant clonal results, which exhibited a T1NB identity but never a T2NB identity (Fig. 4A-E).

We next asked whether SERCA is uniquely required for T2NB identity or whether manipulating other Ca^2+^ regulatory proteins are sufficient to alter T2NB identity. Individual knockdown of *ncx*, *Cam*, *Orai, Stim* or *Itpr* had no discernable effect on T2NB identity, indicating that SERCA is uniquely required among the candidates tested (Supp. Fig. 4A-H). Notably, SERCA is also the only candidate whose knockdown is expected to deplete ER Ca^2+^ while elevating cytosolic Ca^2+^. To distinguish whether SERCA’s regulation of ER luminal or cytosolic Ca^2+^ is more important for T2NB identity, we investigated a double knockdown of *SERCA* and *Orai*. Orai is a plasma membrane Ca²⁺ channel that opens when the ER-localized Ca²⁺ sensor STIM detects ER Ca²⁺ depletion, allowing Ca²⁺ influx that activates SERCA to refill ER stores. Thus, double knockdown of *Orai* and *SERCA* should prevent STIM–Orai–mediated Ca²⁺ entry and reduce the cytosolic Ca²⁺ elevation seen with SERCA knockdown alone ^45^. Double knockdown of *SERCA* and *Orai* nevertheless maintained the T2 to T1NB transformation (Supp. Fig. 4E-E’’’, H), which indicated that the identity change primarily reflected SERCA dependent regulation of ER Ca^2+^, rather than its effect on cytosolic Ca^2+^.

T2NB are specified during embryogenesis ^64,85^ and their identity is maintained until their terminal differentiation during pupation ^74,86^. The continued labeling of *SERCA* RNAi lineages by the T2NB driver (*insc-GAL4; ase-GAL80*) indicates that the lineage was correctly established but lost the ability to maintain its Dpn^+^Ase^-^ identity. To determine the timing of the transformation, we stained larval brains for Dpn and Ase from 24 to 96h ALH. As expected, control T2NB retained their Dpn^+^Ase^-^ identity at all stages examined (Fig. 4J). *SERCA* RNAi T2NBs were largely wild type at 24h ALH but became fully transformed by 72h ALH (Fig. 4J). Although the identity shift clearly occurred at the NB level, SERCA knockdown could also affect downstream progenitors. For instance, the near-complete loss of Dpn^+^ INPs could result from apoptosis. Blocking apoptosis with P35 did not significantly alter INP number in SERCA knockdown T2NB lineages (Supp. Fig. 5A-E), suggesting that INP loss is not due to cell death. Together, these findings demonstrate that SERCA is required to maintain T2NB identity, and that loss of SERCA transforms T2NBs into a “T1-like” fate with subsequent changes to the identity of downstream progeny (Fig. 4K).

### SERCA is required for maintenance of T2NB transcriptional profiles

Previous work has identified a network of transcription factors and signaling pathways required for T2NB identity (Fig. 5A). Epidermal growth factor receptor (EGFR) signaling through its ligand Spitz is essential for T2NB specification from neural epithelial cells during embryogenesis, where it drives PntP1 expression ^85^. Reduction of Trithorax (Trx), Pointed 1 (PntP1), Buttonhead (Btd), Tailless (Tll), or Notch (N) alone are sufficient to induce a T2 to T1NB transformation, indicating that each is required for T2NB maintenance ^60,87–94^. A major driver of this identity switch is ectopic Ase expression, which occurs when any of these genes are reduced; as a result, T2NBs lose the ability to generate INPs and instead produce GMCs directly, like T1NBs ^89,91,95^. Trx, a methyltransferase, maintains T2NB identity and specifies INP progeny by promoting Btd and PntP1 expression ^91^. PntP1 binds a Tll enhancer to activate Tll expression. Tll, in turn, binds to an Ase enhancer repressing Ase expression and thereby preserve T2NB identity ^89,93,95,96^. In T2NB, Notch is transactivated by its ligand Delta in INPs ^92^. Notch activity induces expression of its target genes including Enhancer of Split (E(spl)), which inhibits the transcriptional repressor Earmuff (Erm), permitting PntP1 expression and reinforcing T2NB identity ^88^.

**Figure 5.**
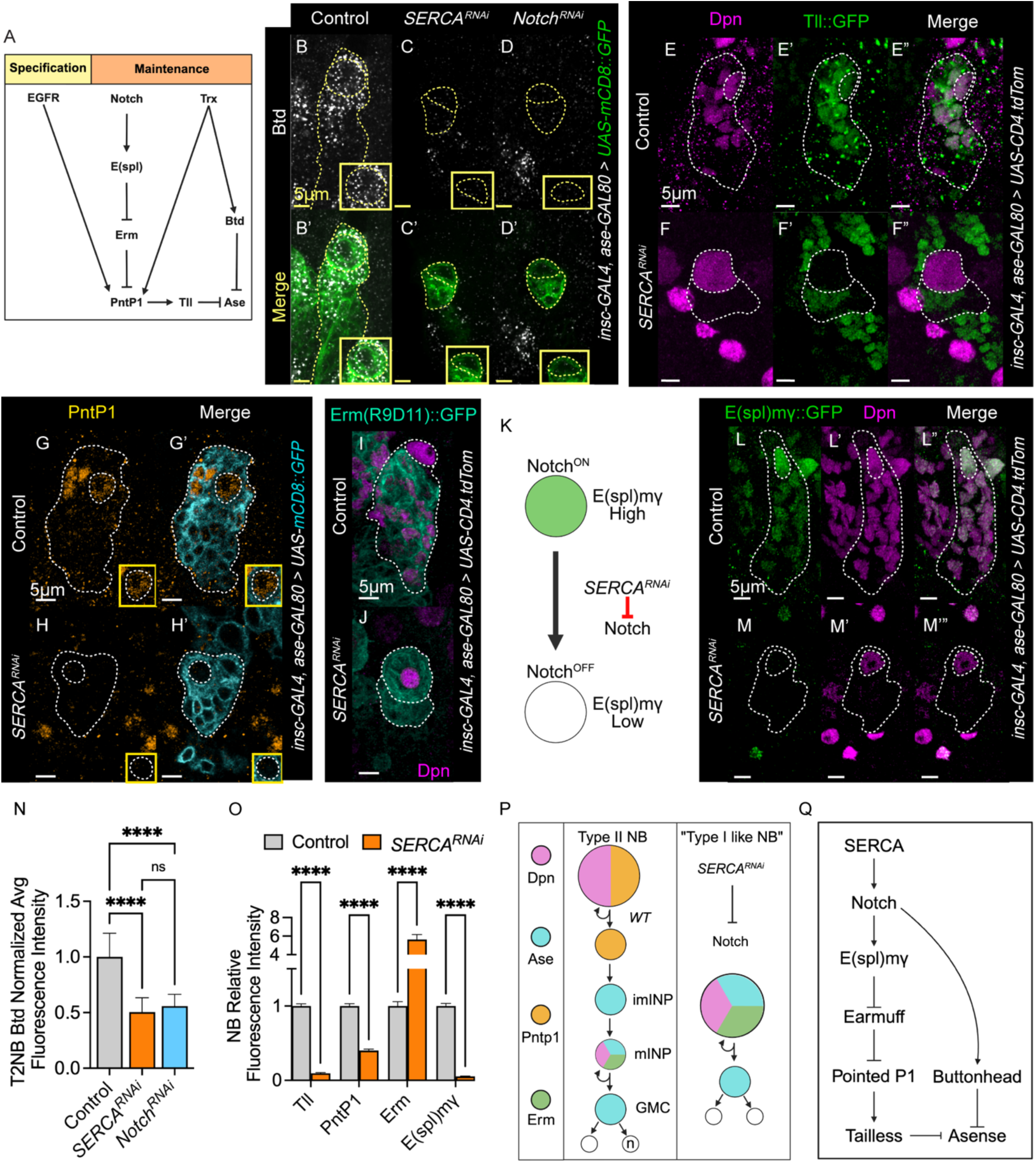
SERCA regulates Notch signaling to maintain type II NB identity. (A) Model of known regulators of T2NB identity. (B-D) Representative images of Buttonhead (Btd) mRNA transcripts in T2NB from third instar larvae. N = 60. (E-F) Representative images of Tailless (*UAS-Tll::GFP*) expression in T2NB from third instar larvae. Lineage outlined using *UAS-CD4::TdTom* (not shown). N ≥ 125 (G-H) Representative images of PointedP1 (PntP1) staining in T2NB from third instar larvae. Lineage outlined using *UAS-mCD8::GFP*. N ≥ 77 (I-J) Representative images of Earmuff (*UAS-Erm(R9D11)::GFP)* expression in T2NB from third instar larvae. Lineage outlined using *UAS-CD4::TdTom* (not shown). N ≥ 44 (K) Illustrative model of *SERCA^RNAi^* effects on Notch activity using *UAS-E(spl)mg::GFP* as a reporter. (L-M) Representative images of Notch activity reporter *(UAS-E(spl)mg::GFP)* expression in T2NB from third instar larvae. Lineage outlined using *UAS-CD4::TdTom* (not shown). N ≥ 107 (N) Quantification of relative fluorescence intensity of control (*luc^RNAi^*, B), *SERCA^RNAi^*(C), and *Notch^RNAi^* (D) Btd transcripts within NB. One-way ANOVA with applied post hoc Tukey’s multiple comparisons test. (O) Quantification of relative fluorescence intensity compared to controls (*luc^RNAi^*) within NB. Independent t-tests. (P) Illustrative model of *SERCA^RNAi^ e*ffect on T2 to “T1-like NB” identity change via Notch. WT (wild type), immature intermediate progenitor (imINP), mature INP (mINP), ganglion mother cell (GMC), neuron (n). (Q) Model of SERCA regulation on known regulators of T2NB identity maintenance. Dashed white lines on representative images outline NBs (circle) and lineages. Yellow inset boxes highlight NB.

Our SERCA transformation timeline reveals that at 24h ALH, most T2NB still resemble wild-type NB and only later ectopically express Ase (Fig. 4J). This indicates that SERCA is required for maintenance rather than initial specification of T2NB identity. To begin dissecting how SERCA supports T2NB maintenance, we first asked whether known regulators of Ase expression were altered.

Btd is required for T2NB identity, including expression and the ability to generate INPs. Ectopic Btd expression in T1NB represses Ase and is sufficient to convert T1NB into T2NB ^91^, indicating that Btd normally functions as an Ase repressor in T2NB. To determine whether *SERCA* knockdown affects Btd, we performed fluorescent *in situ* hybridization and found strong *btd* mRNA expression within control T2NBs (Fig 5B-B’, N). In contrast, *btd* expression was lost upon *SERCA* T2NB lineage knockdown (Fig. 5C-C’, N). Tll is the *Drosophila* ortholog of the orphan nuclear receptor, *TLX (NR2E1, Nuclear Receptor Subfamily 2 Group E Member 1)* and is uniquely expressed in T2NB lineages, where it directly represses Ase by binding upstream of the Ase transcription start site ^93,95^. Loss of Tll could therefore account for the ectopic Ase expression observed upon *SERCA* knockdown. To test this, we used a Tll-EGFP reporter line, in which EGFP is inserted into the endogenous *tll* locus ^93,95,97^. Tll-EGFP was robustly expressed in wild type T2NBs (Fig. 5E-E’’, O) consistent with prior reports ^93–95^, but was drastically reduced following *SERCA* knockdown (Fig. 5F-F’’, O). Because Btd and Tll both normally represses Ase, their loss provides a mechanistic explanation for the ectopic Ase expression in *SERCA-RNAi* animals. Tll expression in T2NB is activated by PntP1 binding to a *tll* enhancer region, so we next asked whether *SERCA* knockdown alters PntP1 expression. T2NB control animals displayed robust PntP1 expression, but was absent upon *SERCA* knockdown (Fig. 5G-H’, O). PntP1 expression is normally present in T2NB but absent in INPs due to Erm repression. Erm is repressed in T2NB by Notch signaling ^88^. Thus, loss of PntP1 in SERCA-depleted animals could reflect dysregulation of Erm or Notch. To explore this possibility, we used an *erm* cis-regulatory reporter fused with membrane-localized GFP ^98,99^. In controls, Erm expression marked INPs but not NBs (Fig. 5I). In contrast, *SERCA* knockdown caused ectopic Erm expression in the T2NB (Fig. 5J, O). Notably, previous work similarly found that Notch knockdown results in ectopic Erm expression in T2NBs ^88^. Together, these findings suggest that SERCA maintains T2NB identity by regulating Erm expression and/or Notch activity.

### SERCA-Mediated Notch Processing Is Essential for T2NB Maintenance and Notch-Driven Tumors

Notch signaling is required for T2NB identity and its loss converts T2NBs into ‘T1-like’ NBs ^60,87,100^. Enhancer of Split (E(spl)) is a Notch target gene and thus a E(spl)-mγGFP reporter has been used widely as a readout of Notch activity (Fig. 5K) ^100,101,102,103^. In control T2NB lineages, E(spl)-mγGFP was strongly expressed, indicating high Notch activity (Fig. 5L-L”, O). In contrast, T2NB knockdown of *SERCA eliminated* E(spl)-mγGFP expression, demonstrating that Notch activity is off or extremely low (Fig. 5M-M”, O). This result supports our hypothesis that SERCA regulates Notch activity. Loss of Notch activity is sufficient to explain the resulting “T1NB-like” identity shifts observed throughout the T2NB lineage upon *SERCA* knockdown (Fig. 5P-Q).

We next tested whether Notch regulates *btd* expression in T2NBs. We found that *Notch* knockdown was sufficient to reduce *btd*, revealing a previously unrecognized Notch–Btd regulatory link (Fig. 5D-D’, O). This evidence indicates that SERCA’s effect on Btd is mediated through Notch (Fig. 7Q). However, we cannot rule out the possibility that SERCA’s regulation is indirect, for example via Trx or other unknown Btd regulatory factors. Together, these results show that key T2NB maintenance factors (Notch, Tll, PntP1, and Btd) are disrupted upon *SERCA* knockdown, driving T2NBs to adopt a “T1NB-like” identity (Fig. 5P).

To test whether SERCA influences Notch activity through its regulation of cytosolic versus ER luminal Ca^2+^, we incubated brains in high extracellular Ca^2+^ for 24h to elevate cytosolic Ca^2+^ and found that this was not sufficient to drive the T2 to T1NB transformation (Supp. Fig 3J-K’’’). Consistent with this, double knockdowns of *SERCA* and *Orai* (which should reduce cytosolic Ca^2+^ levels compared to *SERCA* knockdown alone) did not rescue T2NB identity (Supp. Fig. 4E-E’’’, G-H). As neither manipulation altered T2NB identity this indicated that cytosolic Ca^2+^ levels alone do not explain the T2 to T1NB identity switch. We therefore examined Notch protein levels with an antibody targeting the Notch extracellular domain (N^ECD^). In controls, N^ECD^ was primarily localized to the cell cortex (Fig. 6A-A’, C-D), whereas *SERCA* knockdown caused N^ECD^ to become predominantly internalized (Fig. 6B-B’, C-D). These results suggest that altered Ca²⁺ levels do not simply reduce Notch abundance, but rather, that *SERCA* depletion disrupts Notch receptor processing and trafficking (Fig. 6M).

**Figure 6.**
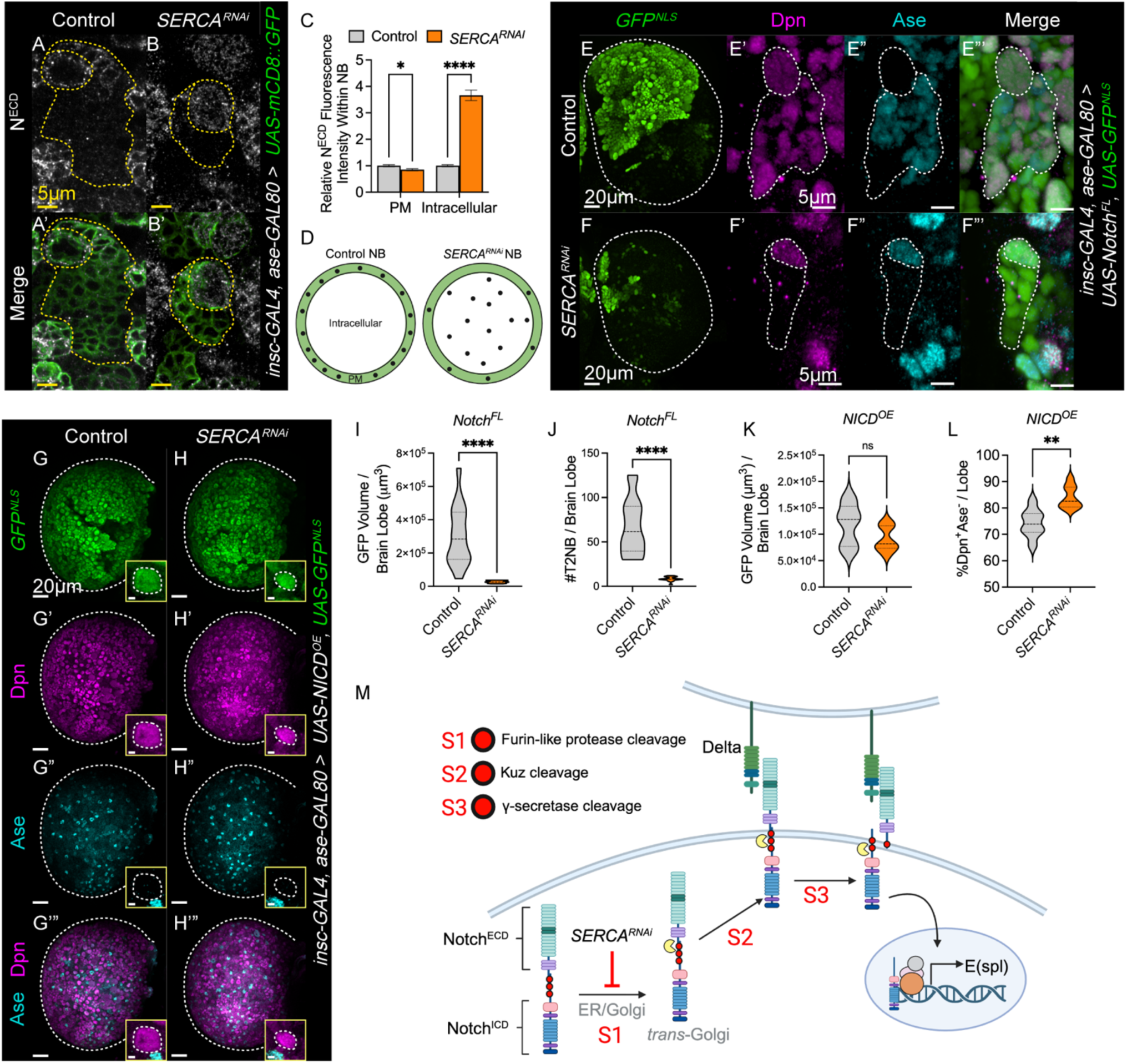
SERCA regulates Notch processing and trafficking to the plasma membrane. (A-B) Representative images of N^ECD^ in control (*luc^RNAi^,* A) and *SERCA^RNAi^* (B) NBs. Yellow dashed line outlines lineage and NB (circle). (C) Quantification of the relative fluorescence intensity of N^ECD^ in control and *SERCA^RNAi^* NBs at the plasma membrane (PM) or intracellular. One-way ANOVA with applied Sidak’s post hoc multiple comparisons test. N ≥ 54. (D) Illustrative model of N^ECD^ localization in (A-B). Green outline = PM. Black dots = N^ECD^. (E-F) Representative images of overexpression of Notch full length (FL) in T2NB with either *luc^RNAi^* (control, E) or *SERCA^RNAi^* (F). Brain lobes outlined in dashed white lines. Zoomed in individual T2NB lineages (E’-F’”) outlined in dashed white lines. 96hALH. (G-H) Representative images of overexpression (OE) of N^ICD^ in control (*luc^RNAi^,* G) and *SERCA^RNAi^* (H) T2NB lineages. Brain lobes outlined in dashed white lines. Yellow inset highlights a single NB (outlined with white dashed line). Scale bar in inset is 2μm. (I-J) Quantification of (E-F) GFP volume per brain lobe (I) and the total number of T2NB per brain lobe (J). Independent t-tests. N ≥ 16 brains. 96hALH. (K-L) Quantification of (G-H) GFP volume per brain lobe (K) and the percent of Ase^-^cells within Dpn^+^ cells per brain lobe (L). Independent t-tests. N = 9 brains. (M) Illustrative model of *SERCA^RNAi^* effects on S1 cleavage of the Notch processing pathway. Yellow Pacman represents S1-3 cleavages.

ER Ca^2+^ is critical for Notch folding and processing. Canonical Notch activation involves sequential cleavages. An initial S1 cleavage in the ER/Golgi by Ca^2+^-dependent Furin-like proteases generates a heterodimeric Notch receptor held together by Ca^2+^-dependent ionic bonds and permits trafficking to the plasma membrane (Fig. 6M) ^104–107^. At the membrane, Delta binding exposes the S2 site for ADAM10/Kuz-mediated cleavage, which in turn exposes the S3 site for γ-secretase–dependent proteolysis, releasing the Notch intracellular domain (N^ICD^) to enter the nucleus and activate target genes such as E(spl) ^108–110^ (Fig. 6M). Consistent with our findings, prior work in the developing Drosophila eye and S2 cells showed that SERCA is required for Notch processing in the ER/Golgi, and that loss of SERCA causes accumulation of uncleaved full-length Notch, indicating impaired S1 cleavage ^106^. Additional work in the *Drosophila* larval wing imaginal disc demonstrated that SERCA is also required for proper intracellular trafficking of Notch and that reducing SERCA expression diminishes Notch signaling ^45^. Thus, our results indicate that ER Ca^2+^ homeostasis maintained by SERCA is required for proper Notch processing and activity in T2NB. Depletion of ER luminal Ca^2+^ by *SERCA* knockdown impairs early processing and trafficking of Notch to the plasma membrane preventing Delta-dependent activation and subsequent N^ICD^ nuclear localization required for Notch activity and maintenance of T2NB identity (Fig. 6M).

Since SERCA loss disrupts early Notch processing, we reasoned that *SERCA* knockdown would prevent tumors driven by full-length Notch (N^FL^) overexpression, but not by N^ICD^ overexpression ^60,69,87,111,112^, which bypasses earlier processing steps. We first overexpressed N^FL^ in T2NB lineages which generated large brain tumors filled with T2NB-like Dpn^+^Ase^-^ tumor cells (Fig. 6E, I-J). Strikingly, *SERCA* knockdown strongly suppressed N^FL^-driven tumor formation and restored approximately eight T2NB per brain lobe (Fig. 6F, I-J). The eight remaining T2NB lineages in the *SERCA* depleted N^FL^ brains contained T1-like Ase^+^ NBs, consistent with our finding that SERCA loss blocks Notch processing required for T2NB identity (Fig. 6F, J). *SERCA* knockdown did not reduce tumor burden when N^ICD^ was overexpressed (Fig. 6G-H, K) and N^ICD^ restored T2-like NB identity in *SERCA* depleted tumors by repressing Ase (Fig. 6H, L). These results indicate that SERCA is required for early Notch processing and trafficking and that *SERCA* depletion is sufficient to suppress N^FL^ driven, but not N^ICD^-driven tumorigenesis (Fig. 6M).

### SERCA expression in T2NB is required for Brat tumor formation

T2NBs normally undergo asymmetric division to generate self-renewing NB and an imINP^58–60^. During NB division, differentiation promoting proteins, Numb and Brain Tumor (Brat) segregate to the basal cortex where they will be inherited by the imINP^59,60^. For newly born immature INPs (imINPs) to mature and commit to their fate, Notch signaling is downregulated ^58–60,99,113^. In imINPs this occurs as Numb antagonizes Notch signaling while Brat promotes mRNA decay of targets that regulate stemness including Dpn and Klumpfuss^113–120^. Together, Brat and Numb ensure that imINPs commit to differentiation. Interestingly, loss of *brat* driven in T2NB lineages (but not T1NB lineages) leads to brain tumor formation (Fig. 7E-F) ^60,114^. Loss of *brat* causes ectopic expression of Dpn and Notch in newly born imINPs, preventing their maturation and reverting them back to a NB-like state, leading to brain tumorigenesis (Fig. 7E) ^112,113,121–123^. Thus, if SERCA transforms T2NBs into ‘T1-like’ NBs, they should be resistant to *brat* knockdown tumor formation. Conversely, if SERCA transforms T2 NB into INP-like cells, they should remain susceptible to *brat*-induced tumors. Strikingly, double knockdown of *SERCA* and *brat* in T2NB lineages prevented tumor formation and instead produced ‘T1-like’ NB lineages (Fig. 7A-D). This outcome demonstrates that SERCA-depleted T2NBs transform into a ‘T1-like’ state that is resistant to Brat-dependent tumorigenesis (Fig. 7F).

**Figure 7.**
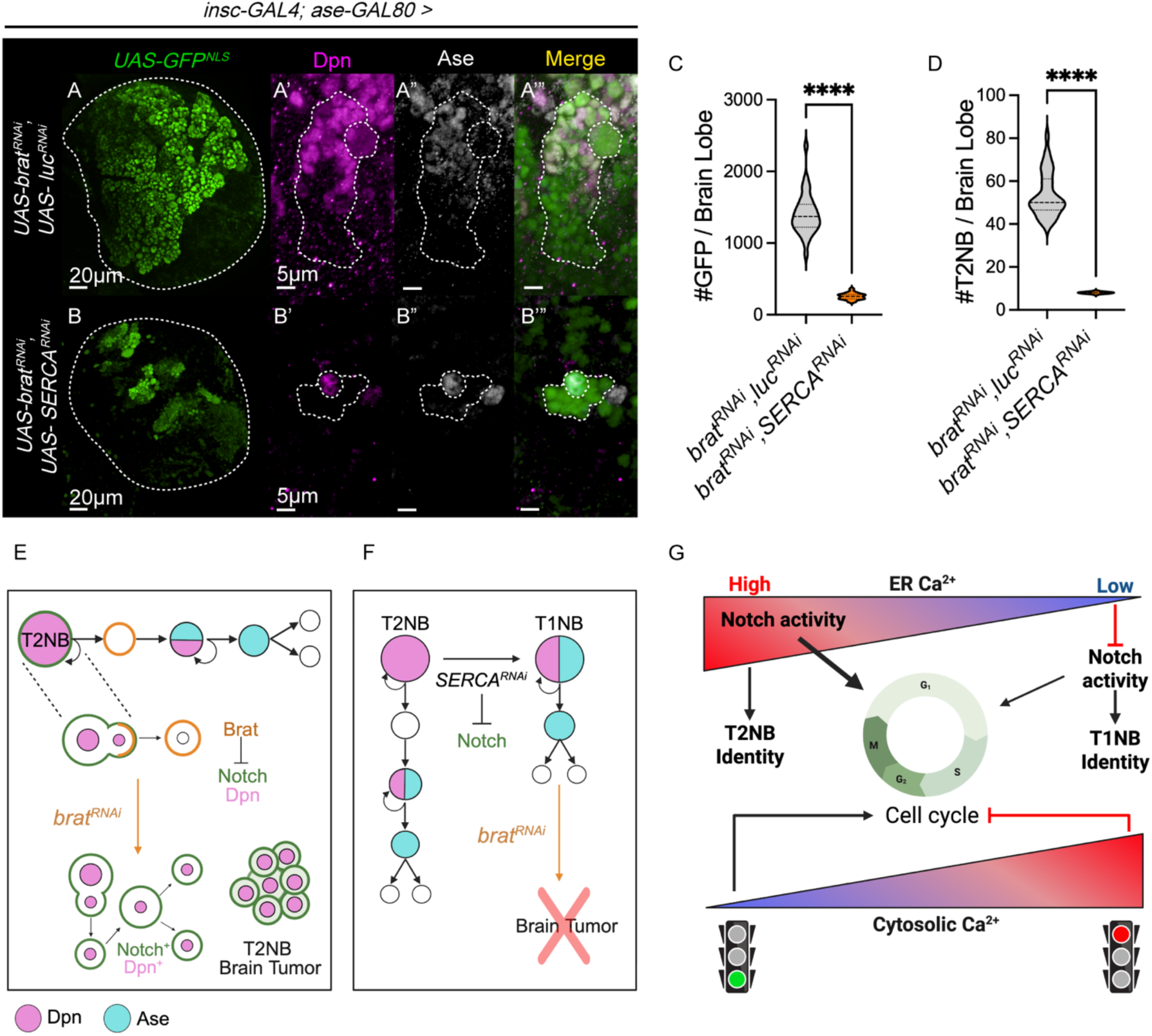
SERCA expression is required for T2NB tumor formation. (A-B) Representative images of control (*brat^RNAi^luc^RNAi^*, A) and *brat^RNAi^SERCA^RNAi^* (B) T2NB brain tumors. Brain lobes (A and B) outlined by dashed white lines. Zoomed in T2NB lineages (A’-B’”) outlined in dashed white lines. (C) Quantification of the number of GFP^+^ cells per brain lobe. Independent t-test. N ≥ 33. (D) Quantification of the total number of T2NB per brain lobe. Independent t-test. N ≥ 33. (E) Illustrative mechanistic model of *brat^RNAi^*^-^induced tumors in T2NB. (F) Illustrative mechanistic model of *SERCA^RNAi^* resistant effects on *brat^RNAi^-*induced T2NB tumors. (G) Summary model of ER and Cytosolic Ca2+ effects on cell cycle progression and NB identity.

## DISCUSSION

Tissue development requires a balance between proliferation and differentiation to generate specialized cell types. In the brain, the production and migration of specific neurons to precise locations is essential for synapse formation and the specificity of neural circuit assembly that underlie information processing. Disruptions in any of these carefully constructed developmental programs can cause developmental disorders. A key question is how a ubiquitous second messenger such as Ca^2+^ can enact distinct physiological outputs across progressively differentiating cell types during brain development. Intracellular Ca^2+^ dynamics are shaped by the expression of ion channels, pumps, and transporters as well as morphological changes that compartmentalize Ca^2+^ domains. These factors generate cell-type-specific Ca^2+^ signaling patterns—spikes, waves, and local “puffs”—that modulate downstream pathways regulating gene expression, cell fate, and behavior. The prevailing view has been that most cells maintain low cytosolic Ca^2+^ with transient increases in response to specific cues.

Our results revise this classic model by showing that baseline Ca^2+^ levels are stratified by cell fate in the developing brain and that progenitors actively modulate this Ca^2+^ landscape to tune canonical developmental signaling pathways and cell behaviors. We find that T2NBs maintain low levels of cytosolic Ca^2+^, which promote proliferation, whereas elevated cytosolic Ca^2+^ acts as a brake that inhibits proliferation and promotes differentiation (Fig. 7G). We also observe significant variation in ER luminal Ca^2+^ across progenitor stages. NBs exhibit relatively low ER luminal Ca^2+^ levels, while INPs/GMCs show higher ER Ca^2+^ levels. These gradients are essential for regulating signaling pathways during differentiation, building on work in fly imaginal discs showing that conserved developmental signaling pathways (Notch, Wingless, and Hippo) differ in their ER Ca^2+^ level requirements ^45^. We demonstrate that luminal ER Ca^2+^ levels are required for Notch processing in T2NBs. Disrupting ER Ca^2+^ homeostasis by knocking down or knocking out *SERCA* expression converts T2NB into a ‘T1-like’ NB fate, and rewires both their transcriptional profile and the identity of cellular progeny, suggesting that dynamic ER Ca^2+^ states and downstream signaling cascades are key drivers of neurogenic fate. Together, our findings support a model in which SERCA-dependent ER Ca^2+^ homeostasis gates Notch processing, thereby coupling baseline Ca^2+^ states to neuroblast identity and tumorigenic potential (Fig. 7G).

Our results share similar Ca^2+^ dynamics with vertebrate systems in the epidermis (skin). Skin stem cells in the basal and spinous layers maintain low Ca²⁺ levels, while more differentiated layers exhibit higher Ca²⁺, forming a Ca²⁺ gradient essential for growth and differentiation^78,124^. Low Ca²⁺ supports self-renewal and division of skin stem cells, while elevated Ca²⁺ promotes their differentiation^125^. In cell culture, increasing extracellular Ca^2+^ is sufficient to induce epidermal differentiation by promoting synthesis of differentiation-specific proteins and regulating cell adhesion molecules that maintain epidermal architecture ^126,127^. The epidermal Ca^2+^ gradient collapses with age, contributing to skin aging phenotypes ^128^. Intracellular Ca^2+^ stores, such as the ER and Golgi apparatus, are major regulators of the skin Ca^2+^ gradient ^129,130^. SERCA-dependent ER Ca²⁺ handling is also critical for neural differentiation. Neuronatin antagonizes SERCA2 to elevate intracellular Ca²⁺, thereby promoting neural induction and differentiation in *Xenopus* and mouse embryonic stem cells ^131^. Like our work, inducing high intracellular Ca^2+^ in mouse embryonic stem cells is sufficient to trigger neural differentiation ^131^. Together with our data, these observations support the idea that tuning baseline cytosolic and ER luminal Ca²⁺ levels is a conserved mechanism for controlling cell fate decisions across tissues and species.

Intriguingly, disruptions in epidermal Ca^2+^ homeostasis are associated with mental health comorbidities. Mutations in human ATP2C1 (encoding the Golgi Ca²⁺ ATPase SPCA1) cause Hailey–Hailey disease, and mutations in ATP2A2 (encoding SERCA2, the closest *Drosophila* SERCA relative) cause Darier’s disease. Both conditions disrupt the epidermal Ca²⁺ gradient and are characterized by blistering skin lesions ^132–134^. Interestingly, patients with Darier’s disease are 4 times more likely to develop bipolar disorder and 2.5 times more likely to develop schizophrenia ^135^. These disorders arise from mutations in Ca²⁺ transporters, and mechanistic work has primarily focused on their effects on postmitotic neural circuitry. Our findings raise the possibility that human SERCA2 also plays important roles in neurogenesis and that SERCA2 mutations may disturb this developmental process, thereby contributing to later-onset neuropsychiatric disorders.

Questions remain regarding whether the *SERCA*-transformed NB more closely resembles a T1NB or an INP, as both NBs and INPs share Dpn^+^Ase^-^ expression profiles. Erm is a regulator of daughter cell differentiation in T2NB lineages ^87,88,112^. In imINPs Erm inhibits T2NB programming as they commit to an INP fate. Maturation of imINPs involves re-expression of Dpn and Notch, followed by new expression of Ase^31,74,136^. Loss of Erm leads to a reversion of the INP back into a T2NB fate when Notch is reactivated^31,94,99,112,137^. Thus it could be argued that the ectopic Erm expression observed in T2NB upon *SERCA* knockdown indicates an ‘INP-like’ identity transformation. However, several lines of evidence argue instead for a ‘T1-like’ NB identity. A recent preprint outlined four criteria to test if T2NB are converted to a T1NB state versus differentiating into an INP-like state^138^. These include the presence of PntP1, Tll, Btd, and expression of T2NB marker Sp1^85,91^. Our results examined 3 of the four criteria and found that Btd, Tll, and PntP1 were all missing (Fig. 5B-G’, R). Thus, we favor the interpretation that SERCA knockdown converts T2NB into a T1NB fate. Ectopic Erm expression may instead reflect a consequence of lost Notch signaling as opposed to a functional readout of INP identity.

In addition, NBs are larger in cell size compared to INPs, which is intimately tied to their identity. Changes to NB size are associated with limits to proliferative potential — this includes quiescence, where NB temporarily exit the cell cycle between embryonic and larval stages, and at the end of their proliferative life where NBs lose the capacity to fully restore volume ^74^. Disturbing factors that drive commitment of imINPs to differentiate results in tumor formation by reverting INPs into a NB fate ^31,87,88,94^. This reversion expands cell size to NB-like levels. Our results found that the SERCA-transformed “T1-like NB” were significantly smaller in NB diameters (but still considerably larger than control INPs). Thus, our data indicates that SERCA knockdown prevents the maturation of NBs into INPs and instead drives them towards a ‘T1-like’ NB fate.

Our manipulations of SERCA influenced tumor formation. Brat is expressed in both T1 and T2NB lineages, but loss of *brat* causes tumors only in T2NBs. This has been attributed to the role of Brat specifically in INPs. Additionally, Btd is required for INP generation within T2NB lineages ^91,96^. Loss of Btd (like our *SERCA* knockdown) transforms T2NB into a “T1-like” NB fate that is resistant to *brat* mutant tumor formation ^91^. This data supports the idea that *SERCA* depletion converts T2NB into a “T1-like” identity, like *trx* and *btd* mutants ^91^. However, an alternative explanation could be that SERCA’s repression of *brat-*knockdown T2NB brain tumors could instead result from the inability of the *brat* INP-like cells to reactivate Notch signaling as we find that loss of *SERCA* represses Notch activity (Fig. 5M). Within NB lineages, Notch is important for self-renewal but also plays a role in neural specification ^139–141^. We found that reduced SERCA expression compromised Notch trafficking and caused T2NB to take on a “T1-like” NB fate. During NB development, daughter cells downregulate Notch which is an important aspect in differentiated progeny as maintaining high Notch throughout NB lineages causes brain tumors. We found that *SERCA* knockdown was sufficient to suppress brain tumors caused by overexpression of full-length Notch, but not those driven by N^ICD^, which bypasses the trafficking steps that depend on SERCA. Because Notch is broadly implicated in tumorigenesis, our results identify SERCA-dependent ER Ca^2+^ handling and Notch processing as potential therapeutic nodes in Notch-driven cancer.

## Acknowledgements

We would like to thank Drs. Jessica Treisman, Yuh-Nung Jan and Cheng-Yu Lee for sharing fly stocks. We would like to thank Dr. James Skeath, Cheng-Yu Lee and Yuh-Nung Jan for sharing antibodies. We would like to thank the Bloomington Drosophila Stock Center for fly stocks and the Developmental Study Hybridoma Bank for Antibodies. We would like to thank Drs. Sijun Zhu, Cheng-Yu Lee, Sarah Certel, Mark Grimes, Jesse Hay and members of the Piggott Lab for comments on the manuscript. This work was supported by grants from the National Institute of General Medicine (P20 GM103546 to Bowler, Pilot grants to B.J.P.), NSF CAREER (NSF 2338239 to B.J.P.), University of Montana startup funding (to B.J.P.). This research was supported by University of Montana Genomics Core and Montana INBRE Data Science Core, which are funded by **the National Institute of General Medical Sciences (P20GM103474)**, the Office of the Vice President for Research and Creative Scholarship at the University of Montana, and the M. J. Murdock Charitable Trust. The content is solely the responsibility of the authors and does not necessarily represent the official views of the UMGC or the National Institutes of Health.

## Materials and Methods

### *Drosophila* stocks and genetics

Flies were reared at 25°C at 70% humidity and grown on standard cornmeal-based food with a standard 12h/12h light/dark cycle. All fly stocks used can be found in Table S1.

To synchronize developmental time points, crosses were placed in bottle cages on 1% grape plates for 8 hours. 24 hours later from the start of transfer to grape plates, plates were completely cleared of L1 larvae. L1 larvae were collected every 2 hours thereafter and transferred to fly food vials. A max of 50 larvae was placed into each vial. Larvae were kept at 25°C until the necessary developmental time points were reached (i.e. 96 hours after larval hatching).

Fluorescent larval balancers were sorted against using a NIGHTSEA Royal Blue (440-460nm excitation) stereo microscope fluorescence adapter. All crosses within the same experiment used the same number of flies to prevent variability due to crowding.

### Immunofluorescence of fixed larval brain tissues

*Drosophila* third instar larval brains were dissected on ice in phosphate-buffered saline (PBS, pH 7.4) for 20 minutes followed by 20-minute fixation in 4% paraformaldehyde (PFA) in PBS followed by three quick washes in PBS and three 20-minute PBS washes on a nutator at room temperature. Samples were then incubated in blocking buffer (2% normal goat serum, 2% Triton-X-100, 2% bovine serum albumin in PBS) for 1 hour at room temperature. Primary antibodies were diluted in blocking buffer and incubated with samples overnight at 4°C (all antibodies used can be found in Table S2). Samples underwent three quick washes in PBS containing 0.3% Triton X-100 (PBST) followed by three 20-minute washes in PBST at room temperature on a nutator before incubation in secondary antibodies for 1 hour at room temperature. To visualize DNA, DAPI (Sigma; Cat no. MBD0015; 1:500) was included in the blocking buffer containing secondary antibodies. Samples were then quickly washed three times in 0.3% PBST followed by two 20-minute washes on a nutator at room temperature and finally one 20-minute wash in PBS. Samples were then cleaned up of excess tissue and mounted in VECTASHIELD (Vector Laboratories; Cat no. H-1900) on glass slides for imaging with vacuum grease on four corners to prevent squishing the tissue with the cover glass. The cover glass was sealed with nail polish on all four sides. All samples were either imaged on a Zeiss 880 Laser Scanning Confocal Microscope or a Nikon Ti2 Eclipse Spinning Disk. Data that compared the fluorescence intensity between samples were all imaged with the same imaging parameters.

### Live imaging of GCaMP in *Drosophila* larval brain tissues

*Drosophila* brains were dissected in sterile Schneider’s medium supplemented with 0.25% human insulin, 1% pen/strep, and 10% FBS then immediately mounted onto microscope slides. Vacuum grease was placed on four corners of slide to prevent the cover glass from squishing the brain. After placing the cover glass on top of the vacuum grease and live brain, supplemented Schneider’s medium was injected under the slide. Nail polish was not used to seal the cover slip to allow airflow. Live brains were imaged immediately after on a Nikon Ti2 Eclipse Spinning Disk Confocal Microscope and excited with a 488nm laser on a 40x water objective with the same imaging parameters for all samples. Samples were never re-imaged to reduce bleaching and phototoxicity effects.

### Ca^2+^ incubation of larval brain tissues

Larval brains were dissected at 72hALH in supplemented Schneider’s medium (0.25% human insulin, 1% pen/strep, 10% FBS) for 20 minutes at room temperature. A portion of the brains were fixed immediately in 4% PFA for 30 minutes. The other portion of brains were incubated for 24h in sterile supplemented Schneider’s medium with either 5mM calcium or 30mM calcium at 25°C with limited agitation. After incubation for 24h, the brains were fixed with 4% PFA for 30 minutes (96hALH). Further immunofluorescence processes remained the same.

### RT-qPCR of SERCA expression

Total RNA was extracted from 20-25 wandering 3^rd^ instar larval brains with TRIzol (Ambion, Ref. No. 15596026). RNA samples were then purified with phenol-chloroform (Invitrogen, Ref. No. 9731G). 1μg total RNA of each sample was used for cDNA synthesis using the High-Capacity cDNA Reverse Transcription Kit (Applied Biosystems, Ref. No. 4368813) following manufacturer’s instructions. qPCR analysis was performed with three biological and three technical replicates with PowerTrack SYBR Green Master Mix (Applied Biosystems, Ref. No. A46109) using a Stratagene Mx3000p Real-time Thermocycler. The results were analyzed using the Pfaffl method. Primers were designed using the UCSC genome browser and SnapGene and purchased through Integrated DNA Technologies. Primer sequences were as follows:

Actin forward: GCG TCG GTC AAT TCA ATC TT
Actin reverse: AAG CTG CAA CCT CTT CGT CA
SERCA forward: GTC CTC TCC ACC GTC TTC CA
SERCA reverse: AAT CTG GGG TTG ACA TCT GCG

### Measurement of fluorescence intensity in NB lineage cells

The average fluorescence pixel intensities were measured using ImageJ. Regions of interest (ROIs) were determined at a single z-stack where the cells of interest were most defined. All genotypes for the same experiment were imaged with the same imaging parameters. Average protein pixel intensities were subtracted by background fluorescence intensities for a final measurement of fluorescence intensity. Relative fluorescence values were acquired by dividing values from the same experiment by the control group’s averages.

### MARCM clone induction

MARCM clones were generated by crossing virgin MARCM-ready flies (BDSC #605351) — containing an *FRT42D* site, *tubp-GAL80*, *FLP recombinase* under control of *tub-GAL4* heat-shock promoter, and *UAS-mCD8::GFP* marker —to male control (FRT42D) or *SERCA^S5^* mutants. 24h ALH larvae underwent heat-shock induction for 1 hour in a 37°C water bath. The developing progeny was returned to 25°C and larval brains were dissected at 96h ALH. Homozygous MARCM clones were labeled with membrane-localized mCD8::GFP, allowing for lineage analysis.

### EdU proliferation assay

All Edu proliferation assays utilized Click-iT Edu Alexa Fluor Imaging Kit (Invitrogen; Cat. No. A10044). EdU and detection reagents in the Click-IT EdU imaging kit were prepared as instructed by the manufacturer. Few granules of bromophenol blue were added to heated fly food and cooled to 50-60°C prior to adding EdU to give a final concentration of 0.2mM. The mixture was poured into plates and allowed to solidify. Larvae were allowed to feed on the EdU-containing medium for 4 hours. Only larvae with blue food visible in their guts were selected for experimentation. Larval brains were dissected immediately after the 4-hour EdU-feeding period then subsequently fixed in 4% PFA.

### Fluorescent *In Situ* Hybridization Chain Reaction (FISH-HCR)

Buttonhead mRNA transcripts were visualized by performing *in situ* HCR using the HCR Gold RNA-FISH kit (Molecular Instruments). *In situ* probes were a generous gift from the Cheng-Yu Lee lab. Wandering third instar larvae were dissected in PBS and fixed in 4% PFA for 25 mins. Fixed samples were washed with PBST containing 0.3% Triton-X-100. Samples were pre-hybridized in hybridization buffer for 45 minutes at 37°C and then incubated with 5nM of primary mRNA Btd HCR probe sets (Molecular Instruments) at 37°C overnight. Samples were washed the following day with HCR HiFi probe wash buffer for 2 x 15 mins at 37°C and 3 x 5 mins at 25°C. Samples were pre-amplified in amplification buffer for 30 mins at 25°C. H1 and H2 secondary hairpins (Molecular Instruments) were denatured at 95°C for 2 mins and promptly snap-cooled on ice for 5 mins prior to addition to samples. 3μM of each secondary hairpin was added to each sample and incubated overnight in the dark at 25°C. Excess hairpins were removed the next day by washing with HCR Gold amplifier wash for 1 x 10 mins then 2 x 30 mins at 25°C. Samples were mounted in ProLong Glass Antifade Mountant (Invitrogen, Ref no. P36982). The Btd mRNA transcripts were imaged using a Nikon Ti2 spinning disk, with the same imaging parameters for all samples done in the same experiment. Buttonhead *in situ* fluorescence intensity was quantified in T2NB using ImageJ.

### Quantification, graphing, and statistical analysis

All images were analyzed and quantified using either Imaris or ImageJ software programs. Data plots and statistical analysis were performed using Graphpad Prism 10 software. *p ≤0.05, **p ≤0.01, ***p ≤0.001, ****p ≤0.0001. “ns” indicates not significant.

---

**Table S1.**
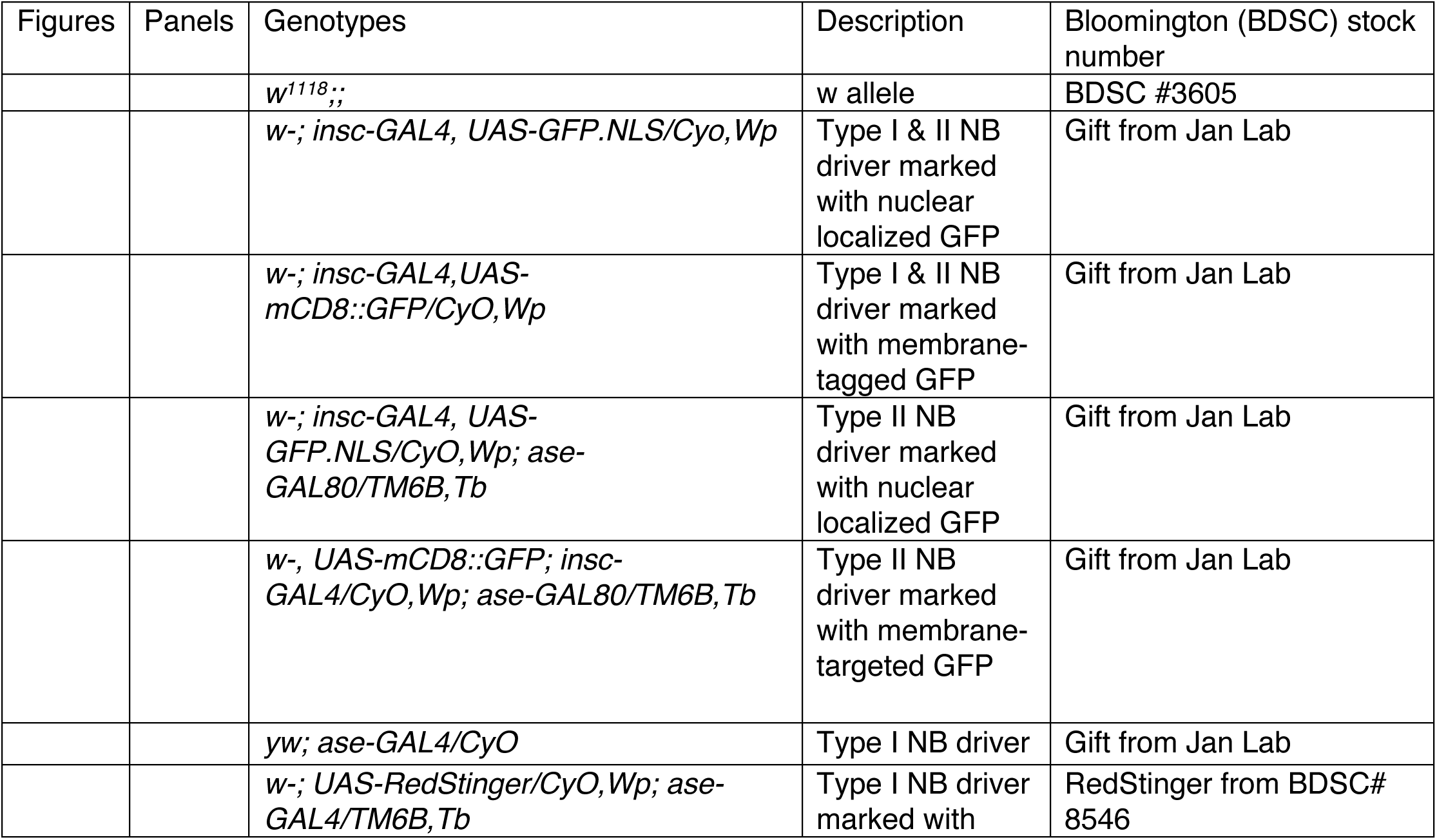
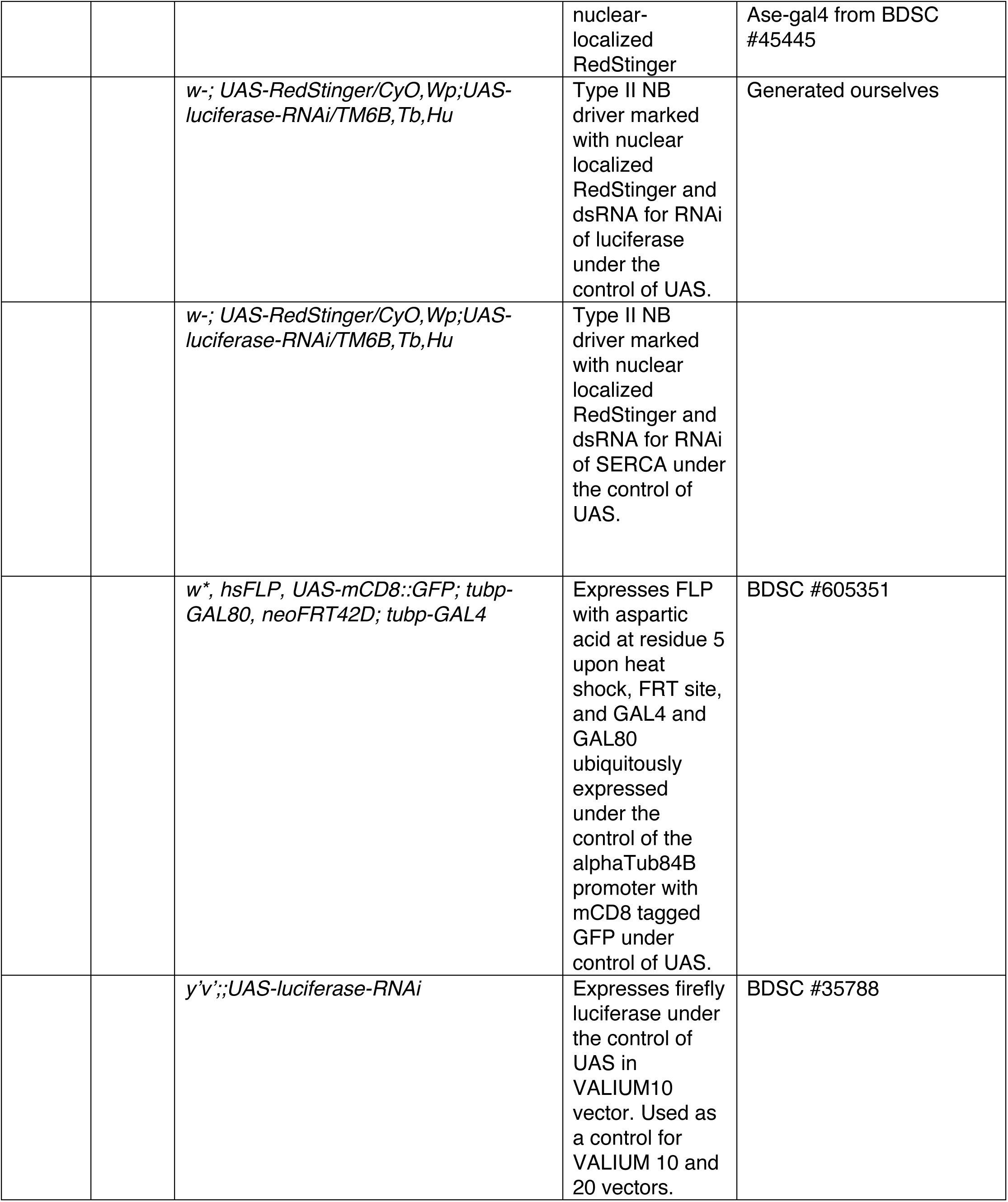
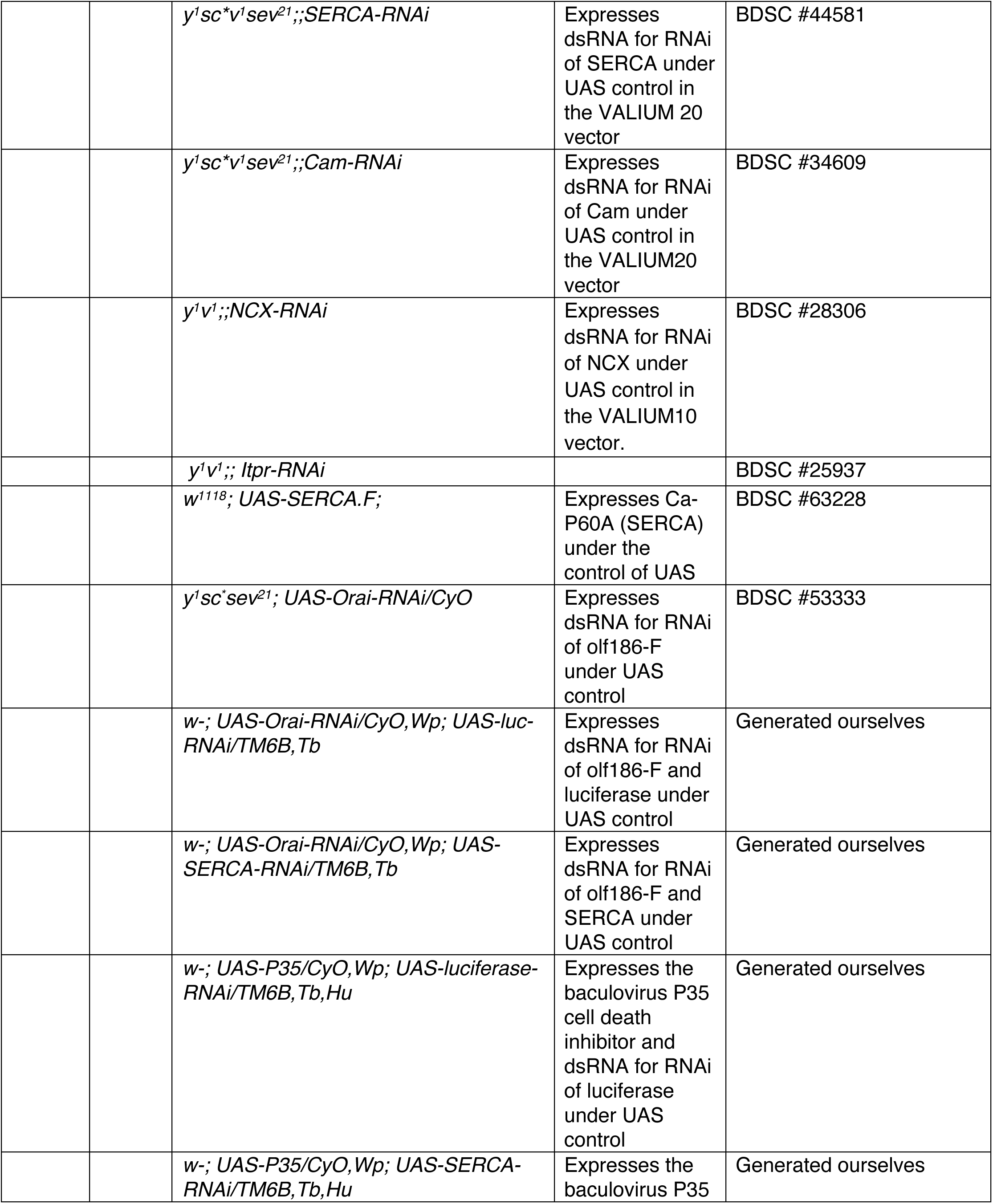
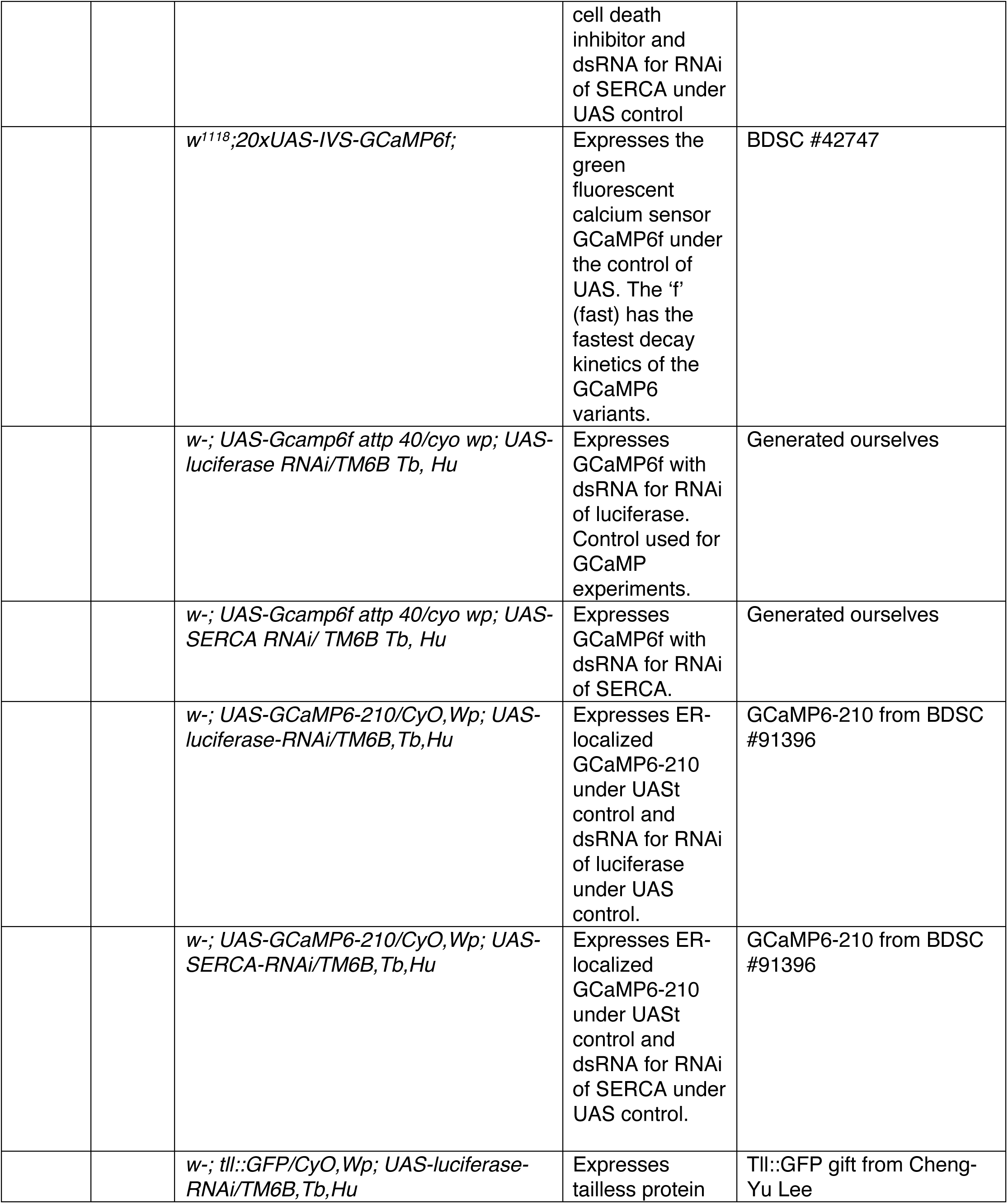
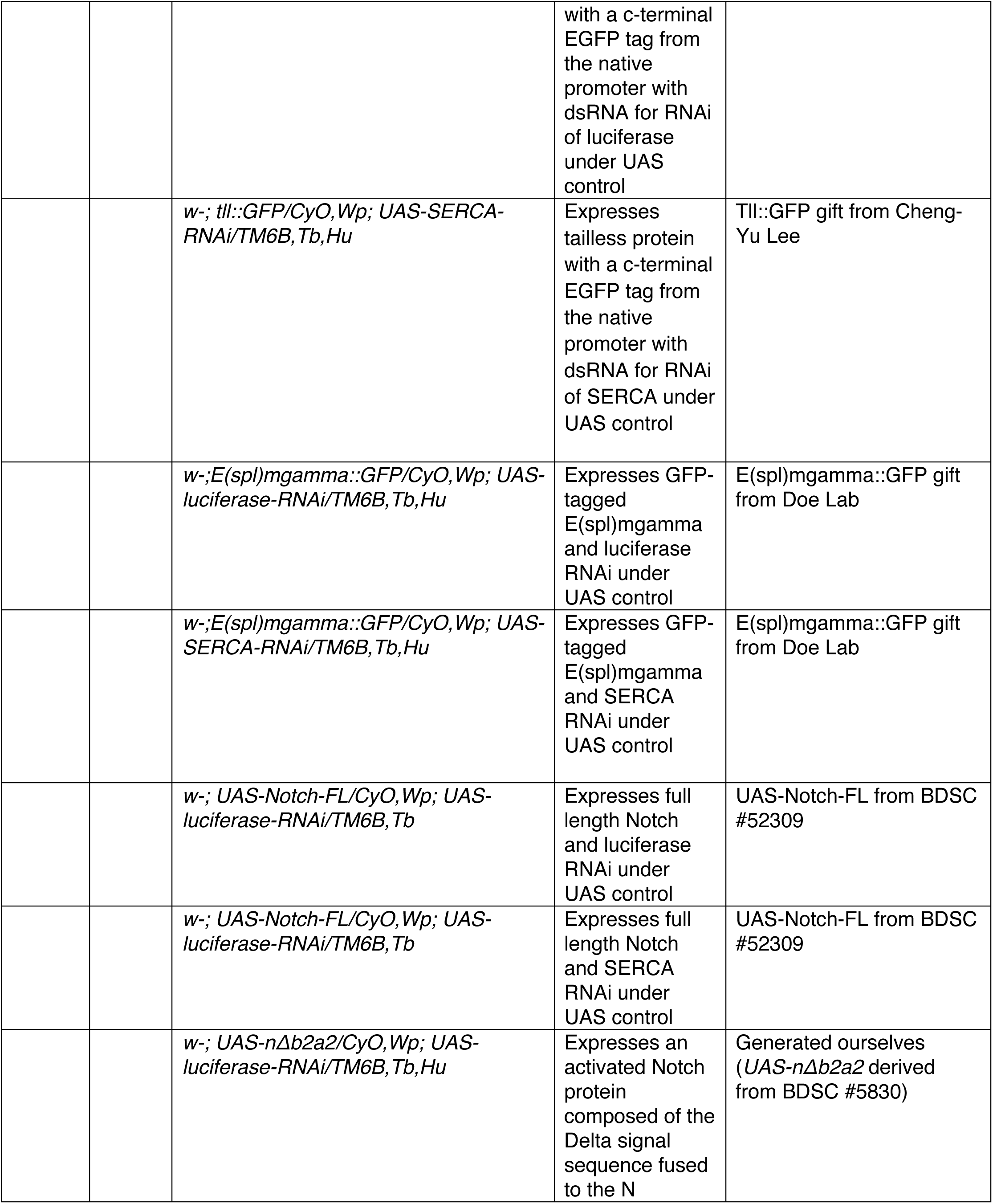
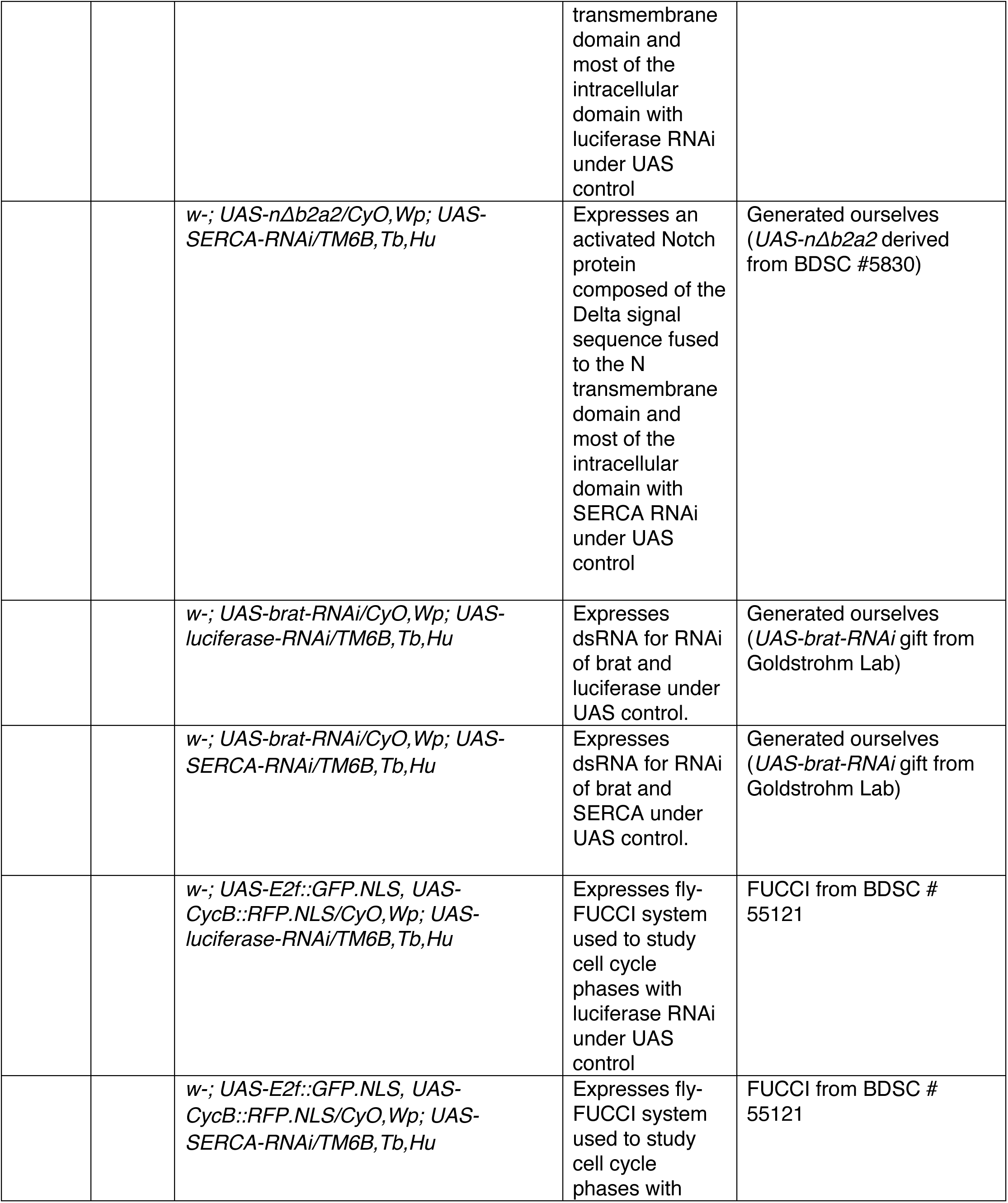
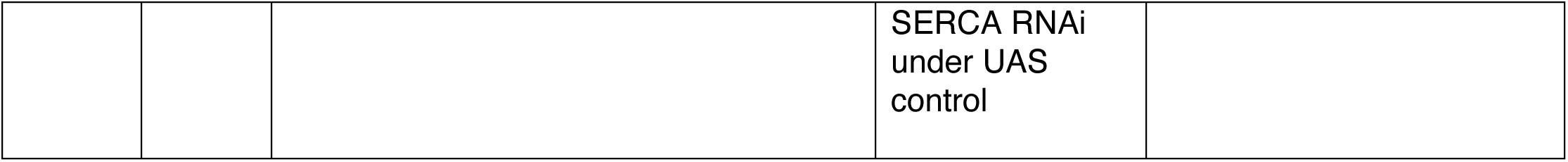
A list of all genotypes for all experimental crosses.

**Table S2.** A list of all antibodies used.

| Figures | Panels | Antibody | Concentration | Source / Cat no. |
| --- | --- | --- | --- | --- |
|  |  | Chicken-anti-GFP | 1:400 | Aves Labs; GFP-1020 |
|  |  | Rabbit-anti-Deadpan | 1:500 | Gift from Yuh-Nung and Lily Jan Lab |
|  |  | Rat-anti-Deadpan | 1:250 | Abcam, ab195173 |
|  |  | Guinea pig-anti-Asense | 1:1000 | Gift from Yuh-Nung and Lily Jan Lab |
|  |  | Rabbit-anti-Asense | 1:400 | Gift from Cheng-Yu Lee |
|  |  | Rabbit-anti-PntP1 | 1:500 | Gift from Skeath Lab |
|  |  | Mouse-anti-Prospero | 1:100 | DSHB; Prospero (MR1a) |
|  |  | Mouse-anti-PH3 | 1:200 | Cell Signaling; 9706S |
|  |  | Rabbit-anti-PH3 | 1:250 | Millipore Sigma; 06-570 |
|  |  | Rabbit –anti-DsRed | 1:500 | Certel Lab |
|  |  | Mouse-anti-NECD (EGF repeats #12-20) | 1:100 | DSHB; C458.2H |
|  |  | Mouse-anti-NECD (EGF repeats #5-7) | 1:100 | DSHB; F461.3B |
|  |  | Rat-anti-TdTomato | 1:500 | Kerafast, EST203 |
|  |  | Goat-anti-Chicken CF488A | 1:400 | Sigma; SAB4600039 |
|  |  | Goat-anti-Guinea Pig CF555 | 1:400 | Sigma; SAB4600064-250UL |
|  |  | Goat-anti-Rat CF568 | 1:400 | Sigma; SAB4600086-250uL |
|  |  | Goat-anti-Mouse CF555 | 1:400 | Sigma; SAB4600067 |
|  |  | Goat-anti-Rabbit<br>CF568 | 1:400 | Sigma; SAB4600085 |
|  |  | Goat-anti-Rabbit<br>CF647 | 1:200 | Sigma; SAB4600185 |
|  |  | Goat-anti-Guinea<br>pig CF640R | 1:200 | Sigma; SAB4600159 |
|  |  | Goat-anti-Rat<br>633 | 1:200 | Sigma; SAB4600142-250uL |
|  |  | DAPI | 1:500 | Sigma; MBD0015 |

**Supplemental Figure 1.**
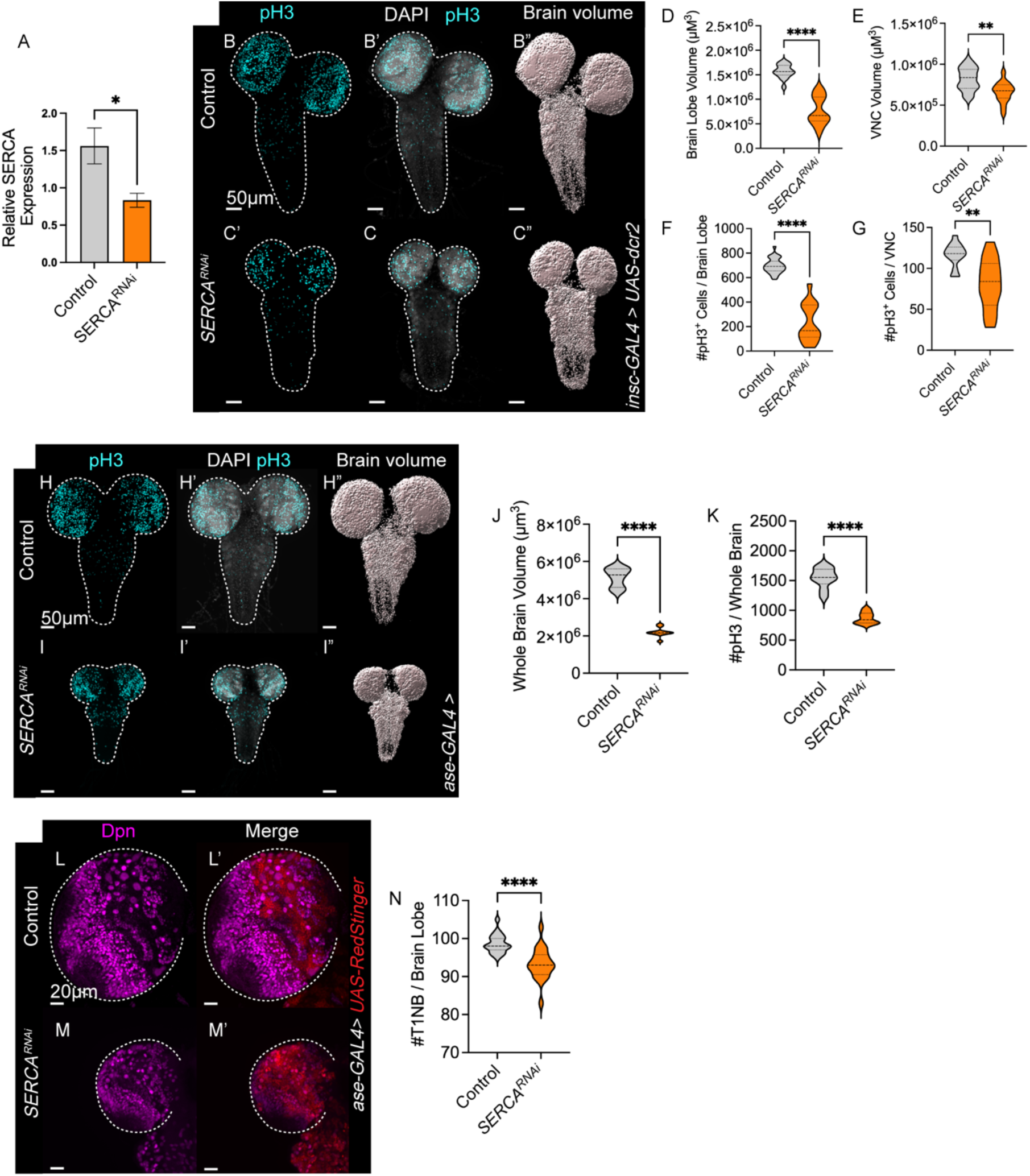
SERCA broadly influences NB lineage development. (A) Quantification of relative SERCA expression in NB from 3rd instar larval brains from real-time qPCR data (pfaffl mathematical model). (B-C) Representative images of control (*luc^RNAi^,* B) and *SERCA^RNAi^* (C) driven in all NB. 96hALH larval brains. White dashed lines outline larval brain. (D-E) Quantification of DAPI volume per brain lobe (D) and per ventral nerve cord (E). Independent t-test. N ≥ 35. (F-G) Quantification of the number of pH3^+^ cells per brain lobe (F) and per ventral nerve cord (G). Independent t-test. N ≥ 35. (H-I) Representative images of control (*luc^RNAi^*, H) and *SERCA^RNAi^* (I) driven in T1NB. 96hALH larval brains. White dashed lines outline larval brain. (J-K) Quantification of (H-I). Whole brain volume (J) and number of pH3^+^ cells per whole brain (K). Independent t-tests. N= 39. (L-M) Representative images of control (*luc^RNAi^*, L) and *SERCA^RNAi^* (M) driven in T1NB. 96hALH larval brain lobes. White dashed lines outline brain lobe. (N) Quantification of (L-M) the total number of T1NB per brain lobe. Independent t-test. N = 20.

**Supplementary Figure 2.**
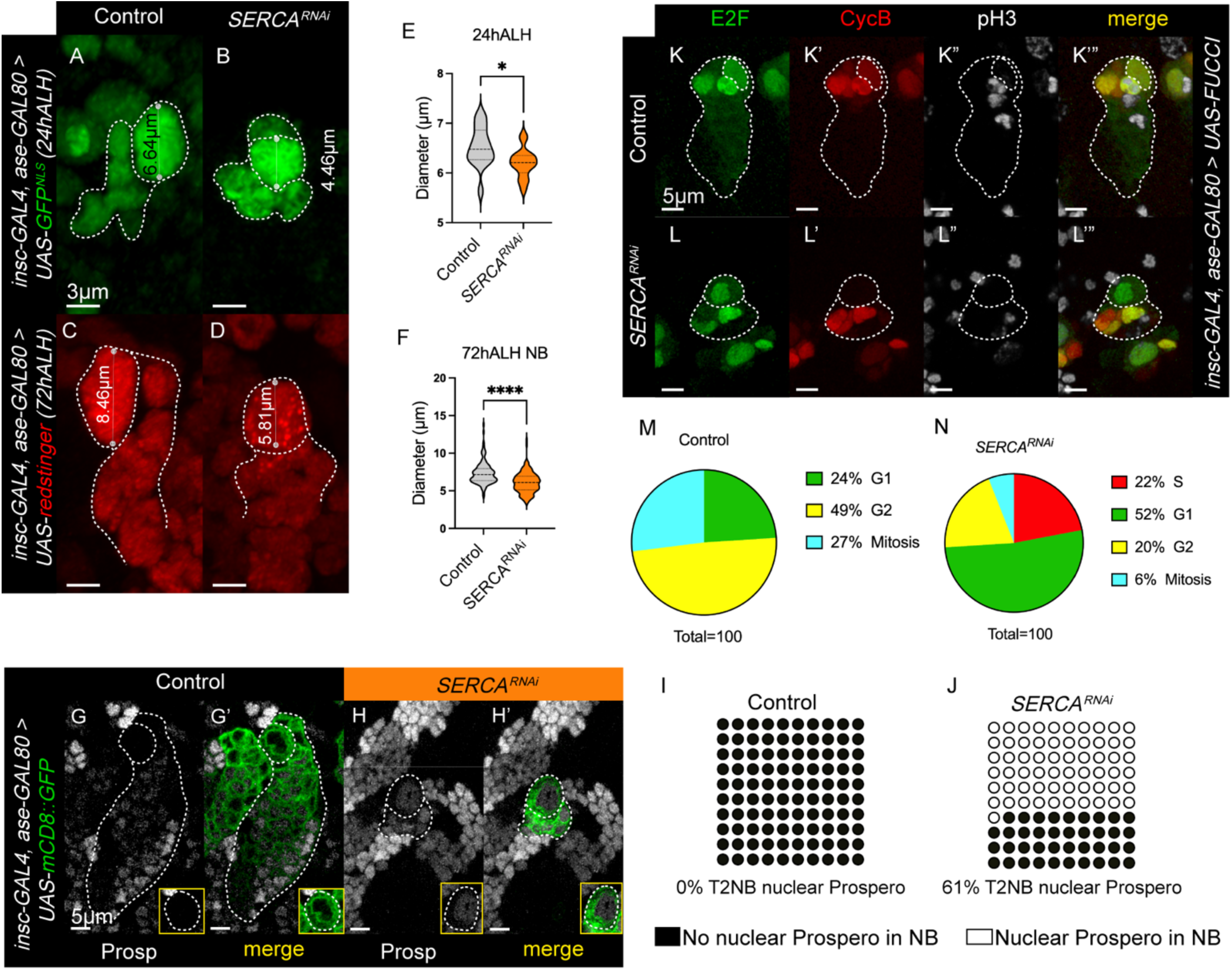
SERCA regulates T2NB cell cycle. (A-B) Representative images of T2NB diameters from 24h ALH control (*luc^RNAi,^* A) and *SERCA^RNAi^* (B) larval brains. (C-D) Representative images of T2NB diameters from 72h ALH control (*luc^RNAi,^* C) and *SERCA^RNAi^* (D) larval brains. (E) Quantification of 24h ALH NB diameters. Independent t-test. N ≥ 16. (F) Quantification of 72hALH NB diameters. Independent t-test. N ≥ 128. (G-H) Representative images of Prospero localization in either control (*luc^RNAi^*, G) or *SERCA^RNAi^* (H) T2NB (labeled with mCD8::GFP). Yellow inset highlights NB. (I-J) Quantification of percentage of T2NB that has nuclear Prospero. Control N= 153 lineages. *SERCA^RNAi^* N = 133 lineages. (K-L) Representative images of FUCCI in either control (*luc^RNAi^*, K) or *SERCA^RNAi^* (L) T2NB. 96hALH. (M-N) Quantification of percentage of cell cycle stages in control (*luc^RNAi^*, M) and *SERCA^RNAi^*, N) T2NB based off FUCCI and phosphohistone 3 staining (pH3). White dashed lines outline NB and lineage.

**Supplemental Figure 3.**
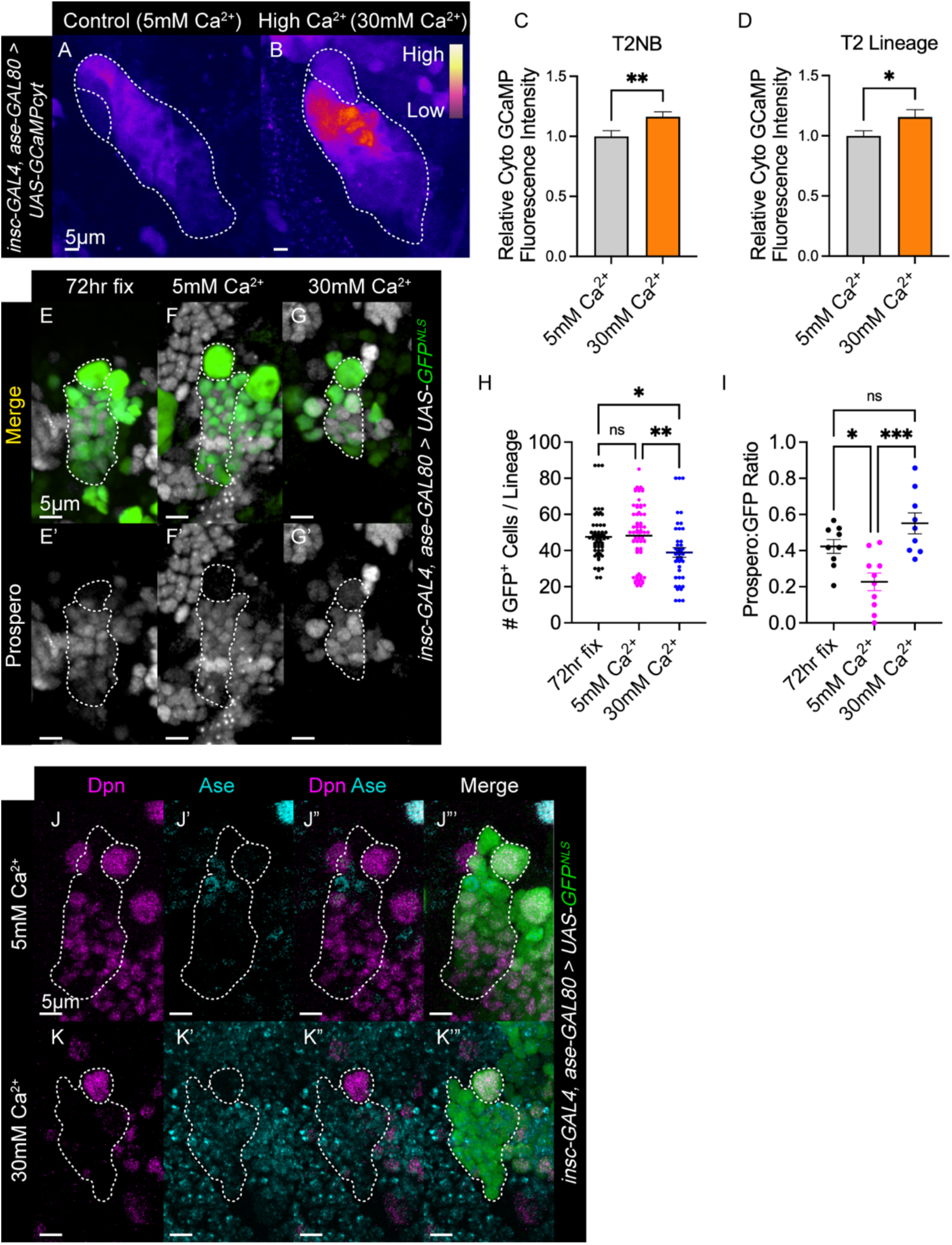
Larval explant brains incubated in high Ca^2+^ medium increased cytosolic Ca^2+^, slowed proliferation, and promoted differentiation. (A-B) Representative images of cytosolic GCaMP (GCaMP6f) expressed in control (5mM Ca^2+^ Schneider’s medium, A) and high Ca^2+^ (30mM Ca^2+^ Schneider’s medium, B) T2NB in 3^rd^ instar larval brains. White dashed lines outline T2NB and lineage. Brighter areas indicate higher Ca^2+^ levels. (C) Quantification of relative cytosolic GCaMP fluorescence intensity in T2NB. Independent t-test. N ≥ 90. (D) Quantification of relative cytosolic GCaMP fluorescence intensity in T2NB lineages. Independent t-test. N ≥ 90. (E-G) Representative images of T2NB of larval brains fixed immediately at 72hALH (E), incubated in 5mM Ca^2+^ (F) for 24 hours, and incubated in 30mM Ca^2+^ (G) for 24 hours. (F-G) dissected at 96hALH after 24h incubation. White dashes outline T2NB and lineage. (H-I) Quantification of the number of cells per T2NB lineage (H) and the ratio of Prospero to *GFP^NLS^* cells per T2NB lineage (I). One-way ANOVA with applied Tukey’s post hoc multiple comparisons test. N ≥ 42. (J-K) Representative images of 72hALH T2NB Ase and Dpn staining in 5mM Ca^2+^ (J) and 30mM Ca^2+^ (K).

**Supplemental Figure 4.**
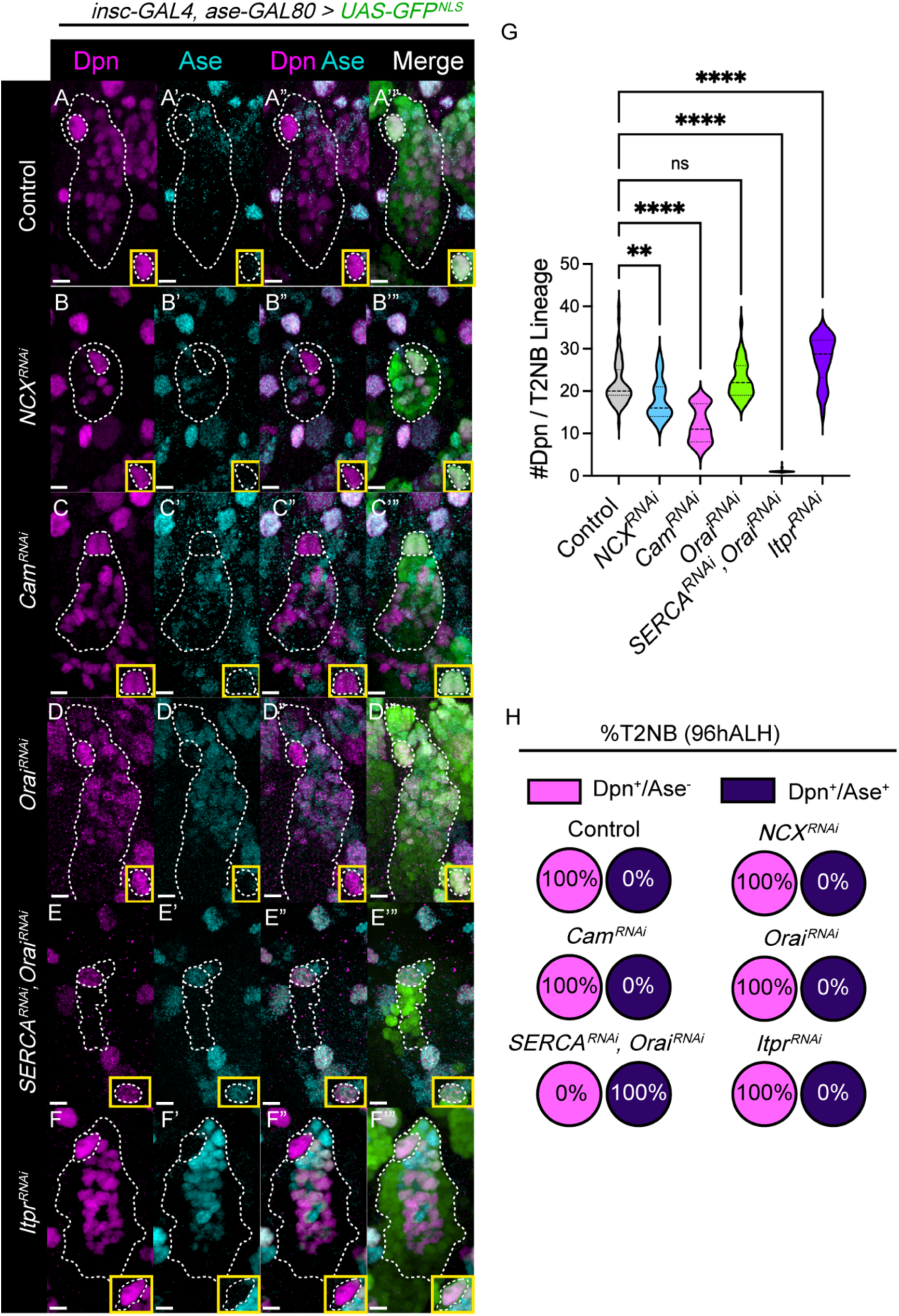
T2NB to T1NB transformation is SERCA-specific. (A-F) Representative images of Ase and Dpn staining in RNAi knockdown of Ca^2+^ regulatory molecules in 96hALH T2NB lineages. Control is *luciferase^RNAi^*. Yellow inset box highlights NB. White dashed lines outline NB and lineage. (G) Quantification of the number of Dpn cells per 96hALH T2NB lineage. One-way ANOVA with applied post hoc Dunnett’s multiple comparisons test. N ≥ 19. (H) Quantification of percentage of T2NB that were Dpn^+^/Ase^-^ or Dpn^+^/Ase^+^. N ≥ 19.

**Supplemental Figure 5.**
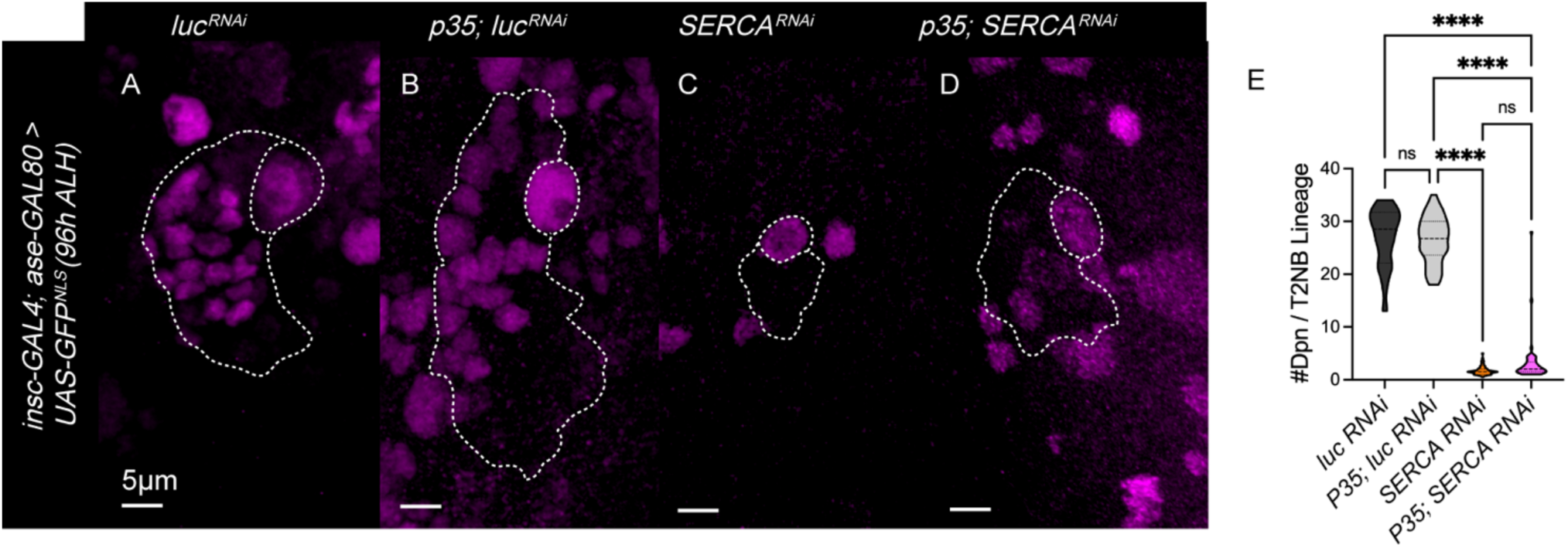
Blocking apoptosis does not rescue INP numbers in SERCA-knockdown T2NB lineages. (A-D) Representative images of RNAi knockdown of control (A) and *SERCA^RNAi^* (C) and RNAi knockdown in conjunction with P35 expression (B, D) in 96hALH T2NB. White dashes outline NB and lineage. (E) Quantification of the number of Dpn cells per T2NB lineage. One-way ANOVA with applied post-hoc Tukey’s multiple comparisons test. N ≥ 20.

